# Estimating uncertainty in family-based GWAS

**DOI:** 10.64898/2026.05.11.724392

**Authors:** Xinyi Miao, Michael D. Edge, Arbel Harpak

**Author notes:** Correspondence should be addressed to A.H.

## Abstract

Standard genome-wide association studies (GWASs) are vulnerable to confounding factors, including stratification, assortative mating, and dynastic effects. Family studies such as sibling-based GWAS (sib-GWAS) mitigate such confounding and are becoming the tool of choice for teasing apart direct genetic effects—causal effects of one’s genotype on one’s own phenotype— from other factors. However, due in part to their smaller sample sizes, sib-GWAS allelic effect estimates are much noisier than standard (i.e., population-based) GWAS estimates. The quantification of this estimation uncertainty is essential for many uses of sib-GWAS, including polygenic scoring, causal inference (e.g., Mendelian randomization), disentangling direct from indirect familial effects, and measuring assortative mating. Here, we investigate sources of uncertainty in sib-GWAS allelic effect estimators. We study their impacts on the biases of three uncertainty measurement methods, including two that are commonly used and a new resampling-based approach we propose. We find that heterogeneity in allelic effects or heteroskedasticity across families (e.g., due to variation in genetic backgrounds or environments) can bias existing methods, and that this bias is more severe for small samples and rare variants. In contrast, the resampling-based approach we propose is approximately unbiased under all scenarios we considered. We validate our theoretical predictions, as well as the importance of effect heterogeneity and heteroskedasticity, using simulations and empirical analysis in the UK Biobank. In sum, this study helps understand the sources of uncertainty in family-based genotype-phenotype association studies and provides a robust method to estimate uncertainty.

## Introduction

Typical genome-wide association studies (GWASs) designs are influenced by population stratification, by assortative mating, and by parental indirect effects^1^. If the goal of the GWAS is to estimate the causal effect of one’s alleles on one’s own phenotype (direct genetic effects)—then these other influences represent confounding. In spite of the risk of confounding, GWAS was adopted by the biomedical research community, largely to the exclusion of family-based study designs such as linkage mapping^2^ and transmission-disequilibrium tests^3^. Although these family-based competitors are much less susceptible to confounding than GWAS, GWAS studies entail much easier recruitment of participants and better statistical power^4^. However, in the last ten years, there has been a resurgence of interest in family-based designs for genetic data^5–7^, driven, among other reasons, by the awareness of residual confounding in standard GWAS^8–12^, and sometimes with the specific motivation of studying indirect familial genetic effects^13–18^, assortative mating^19,20^ or natural selection acting on direct genetic variation^5,21,22^.

One influential family-based design is sibling-based GWAS, sometimes termed sib-GWAS. In a sib-GWAS, the genetic differences among siblings at a given locus are compared with phenotypic differences among them. This design, like its family-based forebears, is largely robust to confounding by population stratification, assortative mating, and dynastic (i.e., parental, grandparental, etc.) indirect genetic effects^23^. Although sib-GWAS-estimated effect sizes are less susceptible to bias than traditional GWAS, they are also less precise, both because typical sample sizes are smaller and also because they are more variable than traditional GWAS even at the same overall sample size.

Despite their importance, there has been little study of the uncertainty of sib-GWAS-based effect estimates. Measures of uncertainty are especially important in analyses of highly polygenic traits, which often integrate signals (and consider signal-to-noise ratio) across numerous sites^24–28^.

Some previous research relied on standard ordinary least squares theory that assumes homoskedastic residuals of allelic effect estimates (henceforth, “OLS method”). However, such an approach will not accommodate heteroskedasticity from heterogeneity in true effect size or from differences among families (sibships) in genetic background or environmental exposures. A second approach, sometimes adopted in light of problems with the OLS method, is to use a permutation distribution to quantify uncertainty^7^. This is the approach used for significance testing in the commonly-used tool, *PLINK*^29^. A recent study suggested a non-parametric approach (clustered standard errors) that is cognizant of possible heterogeneity among families^5^. The accuracy of these measurements of uncertainty has not been evaluated with plausible sources of uncertainty in mind.

There are multiple drivers of uncertainty in allelic effect estimates that require special consideration. First, if true allelic effects are heterogeneous due to gene-by-gene or gene-by-environment interactions that are structured with respect to family membership, then sampling variation will be larger than it would be in the absence of this heterogeneity^30^. Second, variation not explained by the focal locus might be heteroskedastic across families, perhaps due to differences in genetic background or environmental exposures. Third, if an allele (or a binary phenotype) is rare, then there may be few sibling comparisons that are informative about the association.

Here, we focus on the form of sib-GWAS in which there are exactly two individuals in each of *n* sibships (see **Supplementary Material S6** for a generalization to larger families). In this case, the allelic effect can be estimated by regressing the *n* phenotypic differences between sibling pairs on the *n* differences between them in genotype at a focal locus. We consider the previously-used OLS, permutation, and clustered estimators, as well as a new resampling-based method to quantify uncertainty. Specifically, we study the block jackknife^31^, which is more computationally tractable than traditional jackknifing^32^ and more convenient to implement in commonly used tools for GWAS, such as PLINK^29^ and snipar^33^, than bootstrapping^34^.

We mathematically derive the bias of these estimators and validate our results via a simulation study and empirical biobank data. Notably, we show that the OLS variance estimator is biased in the presence of heteroskedasticity, and the permutation variance estimator is biased if the true allelic effect is not zero. Both of these biases are amplified with rare alleles and with small samples. In contrast, the block jackknife variance estimator is approximately unbiased in all situations we studied. Real sib-GWAS data from the UK Biobank^35^ further support our theoretical predictions. We discuss these results in the context of applications of sib-GWAS and recommend the use of resampling-based approaches for accurate assessment of uncertainty in future work.

## Results

### Model

#### Individual-level model

We consider a sibling-based genome-wide association study (sib-GWAS) on a continuous trait. For an individual *j* ∈ {1, 2} in family *i*, the phenotype *Y*_*j,i*_ is modeled as the sum of a family-specific baseline (*Y*_0,*i*_), the additive genetic effect of the focal single nucleotide polymorphism (SNP), and a non-focal term (*ϵ*_*j,i*_):

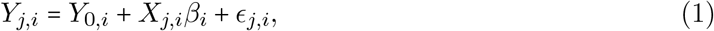

where *Y*_0,*i*_ is the expected trait value when the individual does not carry any effect alleles and *X*_*j,i*_ is the number of reference alleles at the focal site. The non-focal term is assumed to have expectation zero (E [*ϵ*_*j,i*_] =0). We allow the allelic effect *β*_*i*_ to vary among families,

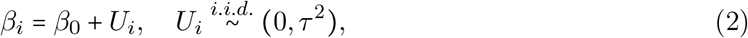

where *β*_0_ is the population-mean effect and *τ*^2^ represents the extent of heterogeneity of allelic effects among families. The non-focal term *ϵ*_*j,i*_ represents the effects of both the environment and genetic background. For most variants influencing complex traits, because each focal SNP typically explains only a minute proportion of phenotypic variance, the variance of the non-focal term is approximately the phenotypic variance. Therefore, across families, variation in within-family non-focal variance should typically correlate strongly with within-family genetic variance.

#### Additive model in sib-GWAS

In a sib-GWAS, we estimate the allelic effect 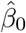 (a local average treatment effect, see Veller, Przeworski, and Coop^30^) by regressing within-family phenotype difference (Δ *Y*_*i*_ = *Y*_1,*i*_ − *Y*_2,*i*_) onto the genotype difference (Δ *X*_*i*_ = *X*_1,*i*_ − *X*_2,*i*_):

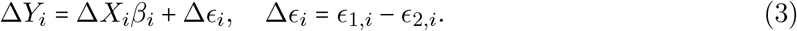

The idea behind this differencing is to mitigate factors that bias allelic effect estimates due to between-family confounding. Importantly, these factors still influence non-focal variance and consequently the variance of allelic effect estimators— our focus of interest in this work. For example, larger focal genotype differences are likelier in families with less assortative mating^19^ and higher levels of heterozygosity in parents^30^. Across families, levels of heterozygosity in parents may proxy a major axis of ancestry, and, in turn, the extent of phenotypic variance^36^ and hence in the extent of non-focal variance. Taken together, this motivates modeling a possible covariance of non-focal variance (Var [Δ *ϵ* _*i*_]) and focal genotype contrast 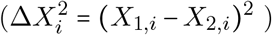:

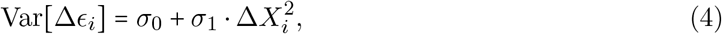

where *σ*_0_, *σ*_1_ are constants. This specification introduces a specific kind of heteroskedasticity in background noise, whereby sibling pairs with greater focal genetic variance (larger 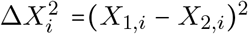) will exhibit systematically higher or lower non-focal variance. While we chose to model heteroskedasticity in a specific, non-exhaustive form, we will later show it appears to capture much of the bias driven by heteroskedasticity in empirical data (see **Empirical sib-GWASs in the UK Biobank**). Together, the heterogeneous allelic effect and the non-focal variation contribute to the uncertainty of the estimated allelic effect in sib-GWAS.

#### Sources of uncertainty are attributable to separable sources

Under the assumptions above, the total variance of the estimated allelic effect 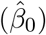 reflects the combined influence of heterogeneous allelic effect (*τ*^2^), the baseline level of non-focal variation (*σ*_0_), and the dependence between non-focal variance and focal genotype contrast (*σ*_1_). The magnitude of the variance is influenced by the sample size and allele frequency of the focal allele. As we show in **Methods**, the uncertainty of the estimated allelic effect is:

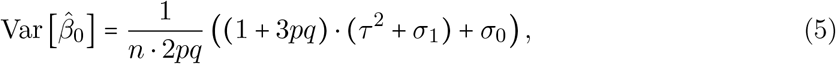

where *p* and *q* = 1 − *p* are the allele frequencies of the reference and alternate alleles, respectively. This key result of our model helps contextualize the separable contribution of these three factors to uncertainty in sib-GWAS.

### Estimation methods

To measure the uncertainty of the estimated allelic effect, we compare three methods: OLS, permutation, and block jackknife. In this section, we will introduce each method and discuss its advantages and biases.

#### The OLS method

As is typical in ordinary least squares (OLS) regression, this method boils down to estimating the variance of 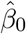 based on the residual sum of squares (RSS) of the fitted linear model **Eq. 3**. Since it corresponds to the OLS regression of within-family phenotype differences on genotype differences, the variance estimator is:

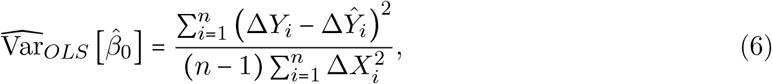

where 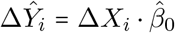 represents the fitted within-family phenotype difference. In this method, the scatter of observed points around the fitted line (**Fig. 1C**) is used to evaluate uncertainty. The OLS variance estimator assumes that the non-focal variation, Δ *ϵ* _*i*_, is homoskedastic and independent of the within-family genotype difference Δ *X*_*i*_.

**Figure 1.**
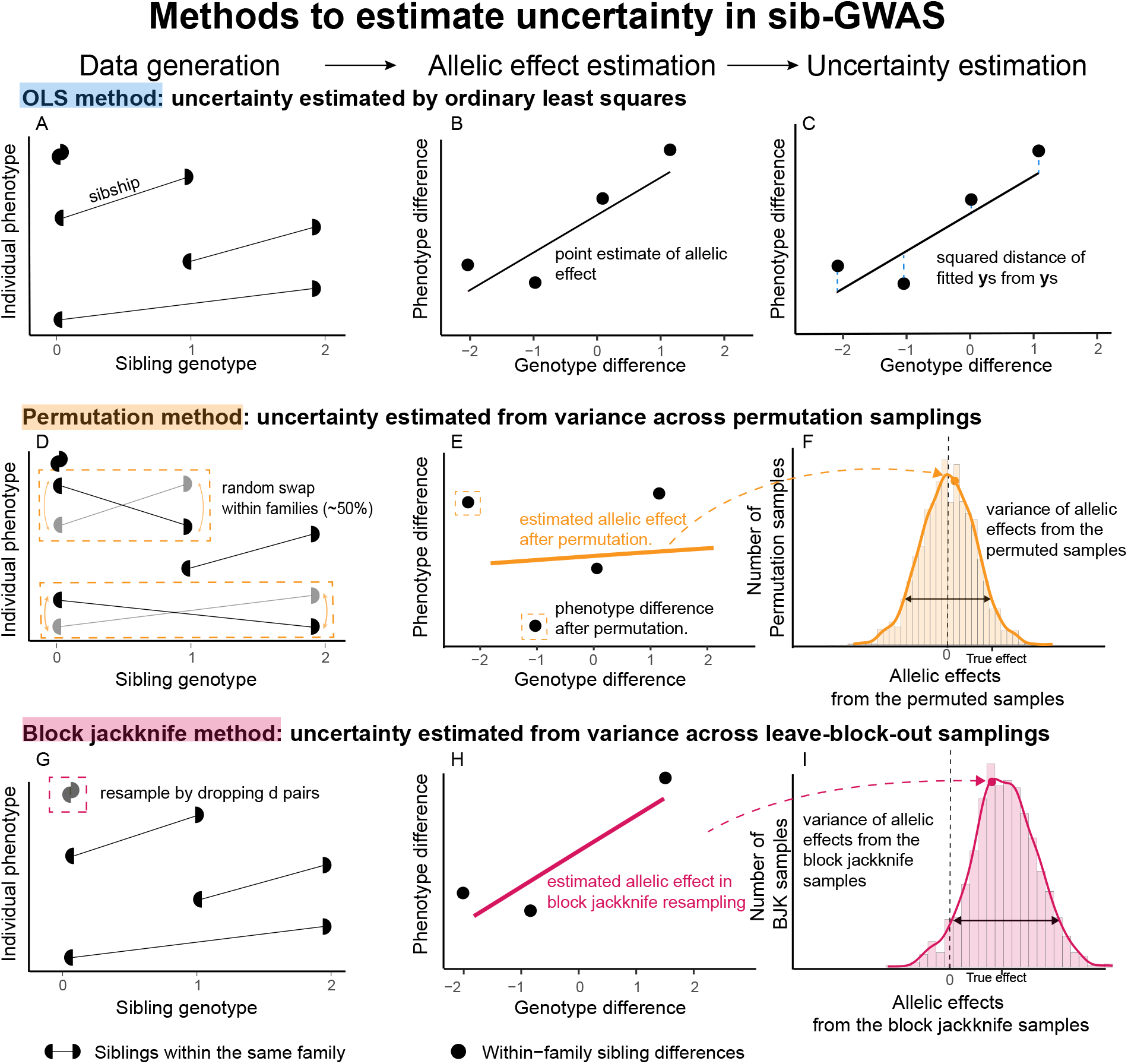
Three methods for estimating uncertainty in sib-GWAS. **(A)** The connected half circles represent siblings from the same family. **(B)** Sib-GWAS estimates the allelic effect by regressing the phenotype difference on the genotype difference. The slope of this regression is the estimated allelic effect. **(C)** The OLS method uses the squared distance between the true value and the fitted value to measure the uncertainty in allelic effect estimates. **(D-F)** The permutation method estimates the uncertainty by generating a null distribution of the allelic effect estimates. The null hypothesis is that there is no true association between genotype difference and phenotype difference. **(D)** The phenotype is swapped at random within sibships, thereby mimicking the null. **(E)** Then the allelic effect is estimated with the permuted data. **(F)** By iteratively repeating the permutation, the re-estimated allelic effects form a null distribution which approximates the variance of allelic effect estimates. **(G-I)** The block jackknife method estimates the uncertainty by iteratively resampling the data. **(G)** First, *d* pairs of siblings are dropped at random. **(H)** Then the allelic effect is estimated with the remaining data. **(I)** By iteratively resampling, the re-estimated allelic effects form a distribution which approximates that of the allelic effect estimator.

Though computationally and analytically straightforward, it can be biased when these assumptions are violated. Specifically, in the **Methods**, we show that the bias of the OLS variance estimator is:

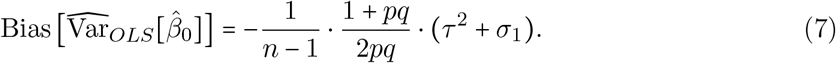

We define the term (*τ*^2^ + *σ*_1_) in **Eq. 7** as the *heteroskedasticity factor* since it captures the combined contributors to bias of two factors that are heterogeneous across families:

1. *τ*^2^ the between-family heterogeneity in allelic effects and
2. *σ*_1_: the slope describing how non-focal variance changes with genotype contrast.

The sign of the combined effect determines the direction of the bias in the OLS variance estimator: a positive heteroskedasticity factor yields a downward bias while a negative factor results in an upward bias (**Fig. 2A**). Additionally, the magnitude of this bias is inversely proportional to the sample size (first term in **Eq. 7**; **Fig. 2C**) and inversely proportional to heterozygosity (second term in **Eq. 7**; **Fig. 2D**).

**Figure 2.**
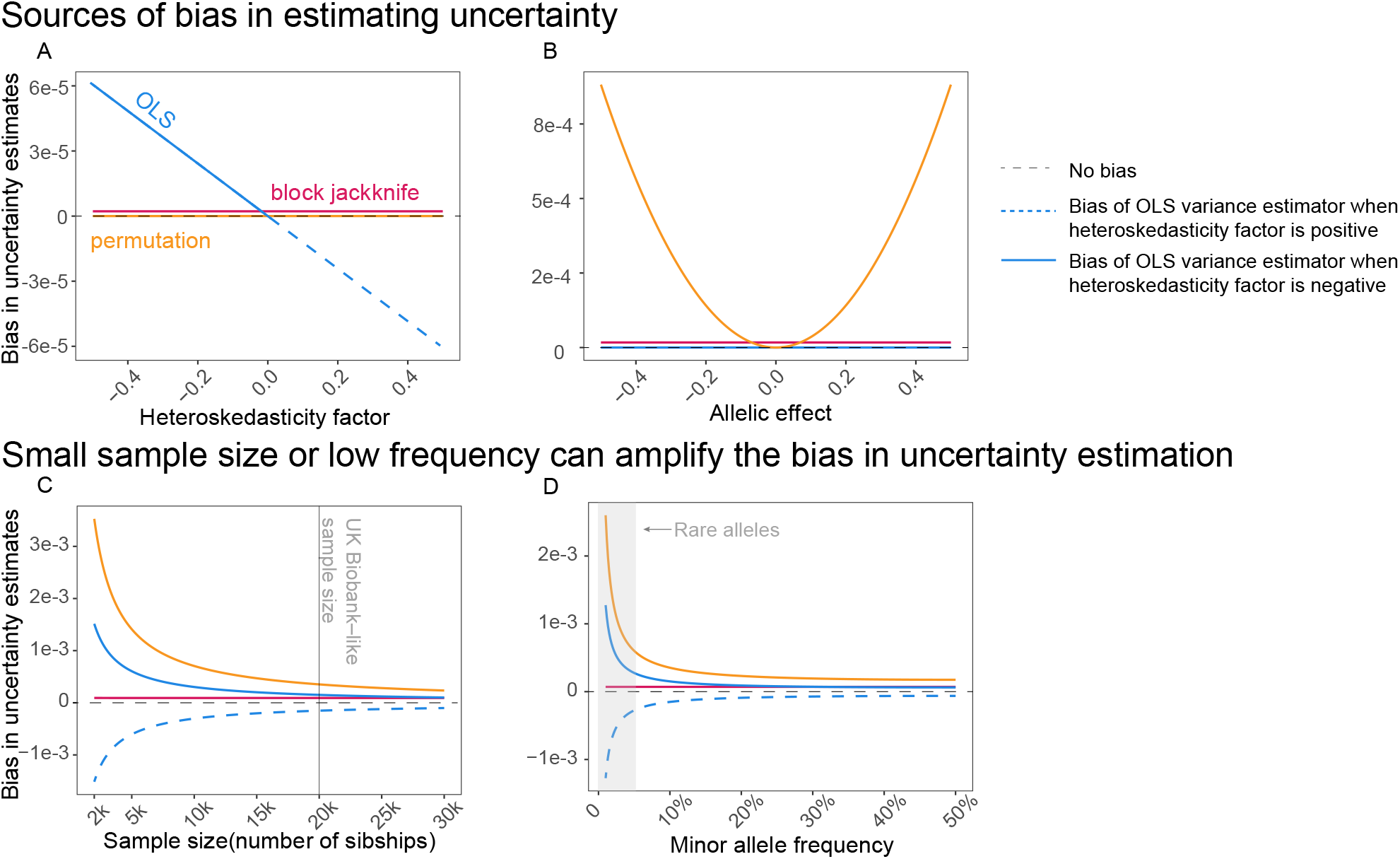
Analytically derived bias of the three variance estimators under varying parameters. The panels show the theoretical predictions made in **Eqs. 7, 8, 9**. Unless otherwise specified, the default parameters are 20,000 sibling pairs and a focal SNP with a maf of 10%, a baseline allelic effect of zero with no variance, and a heteroskedasticity factor of zero. **(A)** varies the heteroskedasticity factor (defined in **Eq. 7**). The OLS variance estimator is biased in the presence of heteroskedasticity. The permutation and block jackknife estimators remain unbiased. **(B)** varies the baseline allelic effect. The permutation variance estimator is upward-biased when the baseline allelic effect is non-zero. The OLS and block jackknife estimators remain unbiased. **(C-D)** vary sample sizes and mafs when the baseline allelic effect is set to 1 for the permutation estimator, and the heteroskedasticity factor is set to −1 (dashed blue line) and 1 (solid blue line) for the OLS estimator. Both the OLS and permutation estimators exhibit large biases when the sample size is limited or the allele frequency is low. In contrast, the block jackknife variance estimator remains approximately unbiased across all scenarios.

#### The permutation method

This approach approximates the uncertainty of 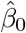 by estimating its uncertainty under the null of no true allelic effect. Namely, an empirical null distribution is generated by randomly permuting sibling phenotypes within families (**Fig. 1(D-F)**). This approach preserves structure across families while breaking the true association within families— between sib differences in phenotype and genotype. For each permuted sample, the allelic effect is re-estimated, and the variance of these permuted estimates provides an empirical measure of the uncertainty of 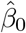. Under the assumption that the allelic effect is zero, the variability across permuted estimates reflects noise in estimation.

Because the permutation method does not rely on the assumption of homoskedastic non-focal variation, the permutation variance estimator is robust to both heterogeneous allelic effect and dependence between non-focal variance and genotype contrast. However, when the true allelic effect is non-zero, the permutation variance estimator is biased (**Fig. 2B**). In **Methods**, we show that

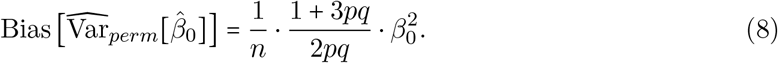

The permutation variance estimate becomes upward-biased in the presence of nonzero allelic effects. In the **Supplementary Material ??**, we show that, in scenarios with more than 2 siblings per family, the bias becomes negative when the allelic effects are heterogeneous (Var [*β*_*i*_] >0). Additionally, the magnitude of the bias is proportional to the squared true allelic effect (**Fig. 2B**), and, as with the OLS variance estimator, it is inversely proportional to sample size and heterozygosity (**Fig. 2(C-D**)).

#### The block jackknife method

As an alternative, we propose a new uncertainty estimator: a block jackknife-based variance estimator, wherein the uncertainty of 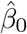 is estimated based on an approximation of the empirical distribution of the allelic effect estimates generated via subsamples of the data (**Fig. 1(G-I)**). In each jackknife replicate, sibling pairs are omitted at random from the dataset (**Fig. 1G**), and the allelic effect is re-estimated using the remaining data (**Fig. 1H**). After multiple jackknife replicates, the variance of these subset-based estimates provides an empirical measure of the uncertainty of 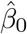, with variability across iterations reflecting sampling variation (**Fig. 1I**).

Similar to the permutation method, the jackknife strategy does not rely on the assumption of homoskedastic non-focal variation; it is robust to heterogeneous allelic effect and dependence between non-focal variance and genotype contrast. Furthermore, the jackknife strategy does not assume a null hypothesis of *β*_0_ = 0 and therefore, as we will show in **Methods**, remains asymptotically unbiased. The traditional jackknife^32^ requires building *n* leave-one-out subsamples, one for each family, and estimating the allelic effect repeatedly across all of them. This exhaustive resampling is computationally intensive for genome-wide studies involving thousands of families. To address this limitation, we adopt the block jackknife method, which omits multiple sibling pairs (“blocks”) in each replicate instead of single observations. This block design substantially improves computational efficiency at the cost of a small penalty in accuracy.

With a limited sample size and resampling replicates, we prove in the **Methods** that the bias of the block jackknife variance estimator is:

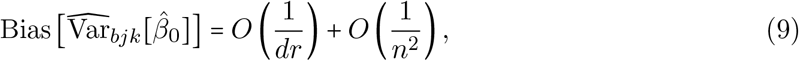

where *d* is the block size (the number of dropped sibships) and *r* = *n* − *d* is the number of remaining sibships in each resampling (**Fig. 2**). Though the theoretical bias of the block jackknife variance estimator is negligible, its empirical stability depends on the number of resampling replicates. Insufficient replicates can inflate noise in the estimated variance. An excessive number of replicates can make computation unnecessarily intensive. To ensure stable uncertainty estimation, the number of replicates must exceed a minimal threshold that balances computational cost and precision. In **Methods**, we follow this rationale and suggest a way to choose the number of block jackknife samplings. Specifically, to ensure that the estimated standard error is within a *δ* − fraction of the true value with probability at least 1 − *α*, it suffices to choose the replicates *m* such that:

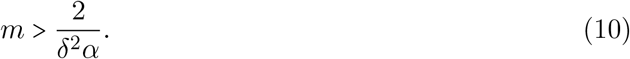

For example, if we wish for the estimated standard error of the allelic effect estimate to be off from the true values by up to 20% with 90% probability, then we would need 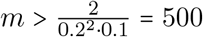 block jackknife resampling replicates.

### Empirical sib-GWASs in the UK Biobank

We sought to evaluate our theoretical results regarding the biases of the three variance estimators using empirical data from the UK Biobank. Because the block jackknife variance estimator is asymptotically unbiased, we used it as an empirical benchmark. Specifically, we calculated the three variance estimates using sib-GWASs we performed in the UK Biobank for 11 physiological traits and 6 social or behavioral traits in a sample of 17,353 sibling pairs. For each SNP, we calculated the ratio of the permutation (or OLS) variance estimate to the block jackknife variance estimate. Ratios greater than 1 are consistent with upward bias, and ratios less than 1 are consistent with downward bias, given that the block jackknife is asymptotically unbiased.

We first tested the prediction of **Eq. 8**, wherein the permutation variance estimator is upward-biased for SNPs with non-zero allelic effects. Since we do not know the true allelic effects, we used SNPs’ sib-GWAS permutation-based *p*-value as a measure of evidence of a non-zero effect. As we predicted based on our derivation, the magnitude of the discrepancy between the permutation and block jackknife variance estimates increased as the evidence of a nonzero effect strengthened (**Figs. 3A, S7**). Additionally, as suggested by the second term of **Eq. 8**, the discrepancy between the permutation and block jackknife variance estimates was larger for rarer alleles (**Figs. 3C, S9**, right-hand panel). Though the permutation variance estimator is expected to have non-negative bias, the ratios fell below 1 for rare SNPs with large *p*-values (> 0.05; **Figs. 3C, S9**, left-hand panel). However, as we explain in **Section S4.2**, this is a parametric regime where this approach for estimating the bias of the permutation estimator is in itself heavily biased.

**Figure 3.**
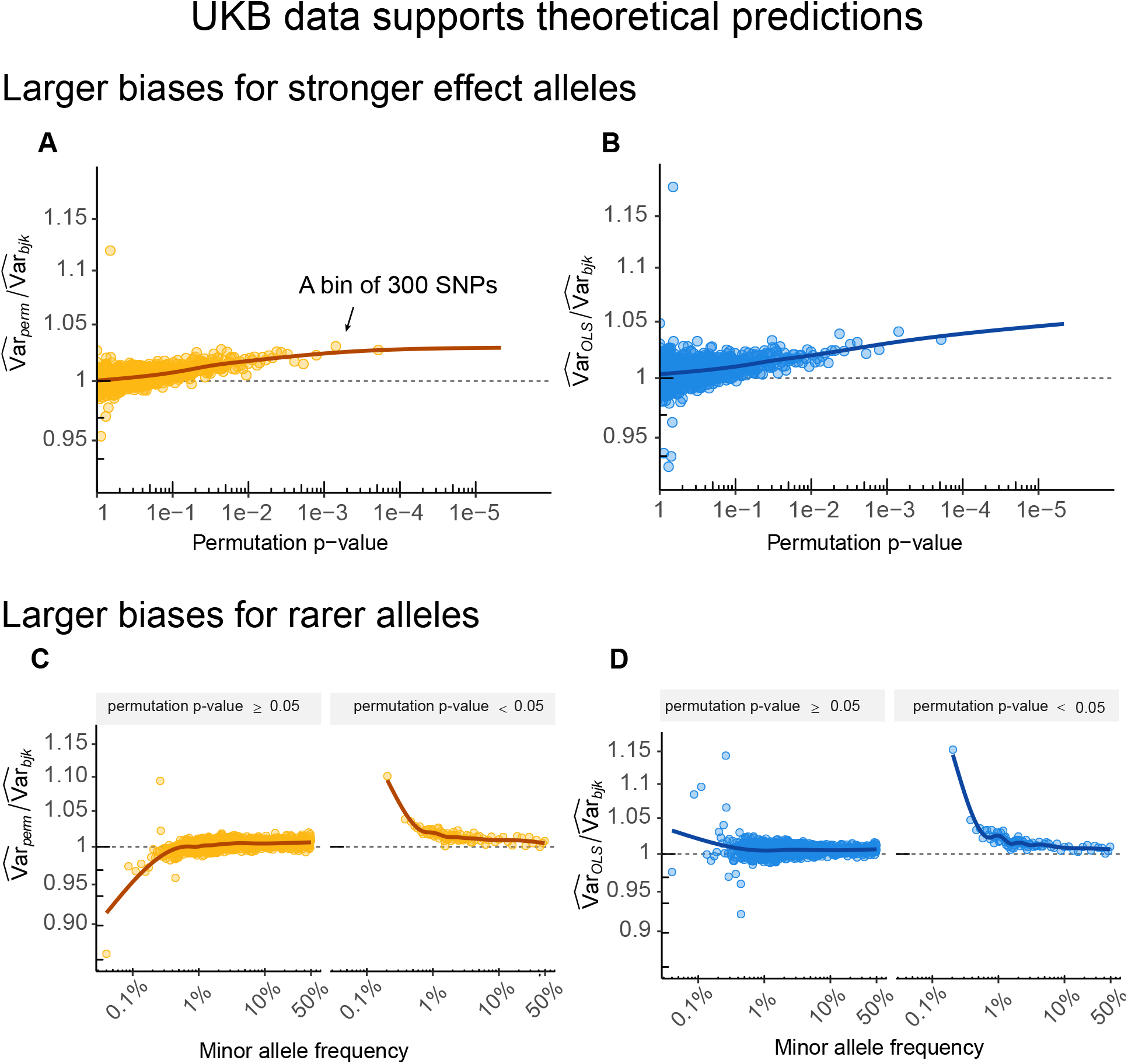
Empirical validation of theoretical predictions in the UK Biobank. To evaluate our analytical predictions in **Eqs. 7, 8**, we performed sib-GWAS for diastolic blood pressure in the UK Biobank using 504,858 SNPs in total. Each point represents averages across 300 SNPs, binned by either the *p*-value of the allelic effect estimate (A-B) or by minor allele frequency (C-D). The x-axis shows the mean value of SNPs in each bin, and the y-axis shows the mean log_2_ ratio of each variance estimator to the block jackknife benchmark. Lines show cubic splines fitted over the raw (unbinned) ratios. **(A-B)** For SNPs with stronger evidence for nonzero allelic effects (smaller *p*-values), both the OLS and permutation estimators deviate from the block jackknife benchmark. The permutation estimator increasingly overestimates variance as evidence for nonzero allelic effects strengthens, consistent with our theoretical expectations. **(C-D)** The discrepancies between the OLS (or permutation) and block jackknife variance estimates are larger in magnitude for rarer alleles. Results for 16 other traits are shown in **Figs. S9, S10**.

We next tested the prediction of **Eq. 7** that the OLS variance estimator is biased downward if the non-focal variance increases with the focal genotype contrast 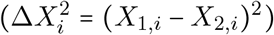 and biased upward when the relationship is reversed. We estimated 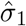, the slope of non-focal variance regressed on the focal genotype contrast for each SNP (see **Methods**). We then used the ratio between the OLS variance estimate and the block jackknife estimate to quantify the discrepancy between the OLS estimator and the block jackknife benchmark across SNPs. As predicted, the ratio was smaller than 1 when 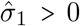 and greater than 1 when 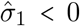 (**Figs. 4, S11**). As with the permutation estimator, the discrepancy between the OLS and block jackknife variance estimates was larger in magnitude for SNPs with smaller *p*-values (**Fig. 3B, S8**) or rarer SNPs (**Fig. 3D, S10**).

**Figure 4.**
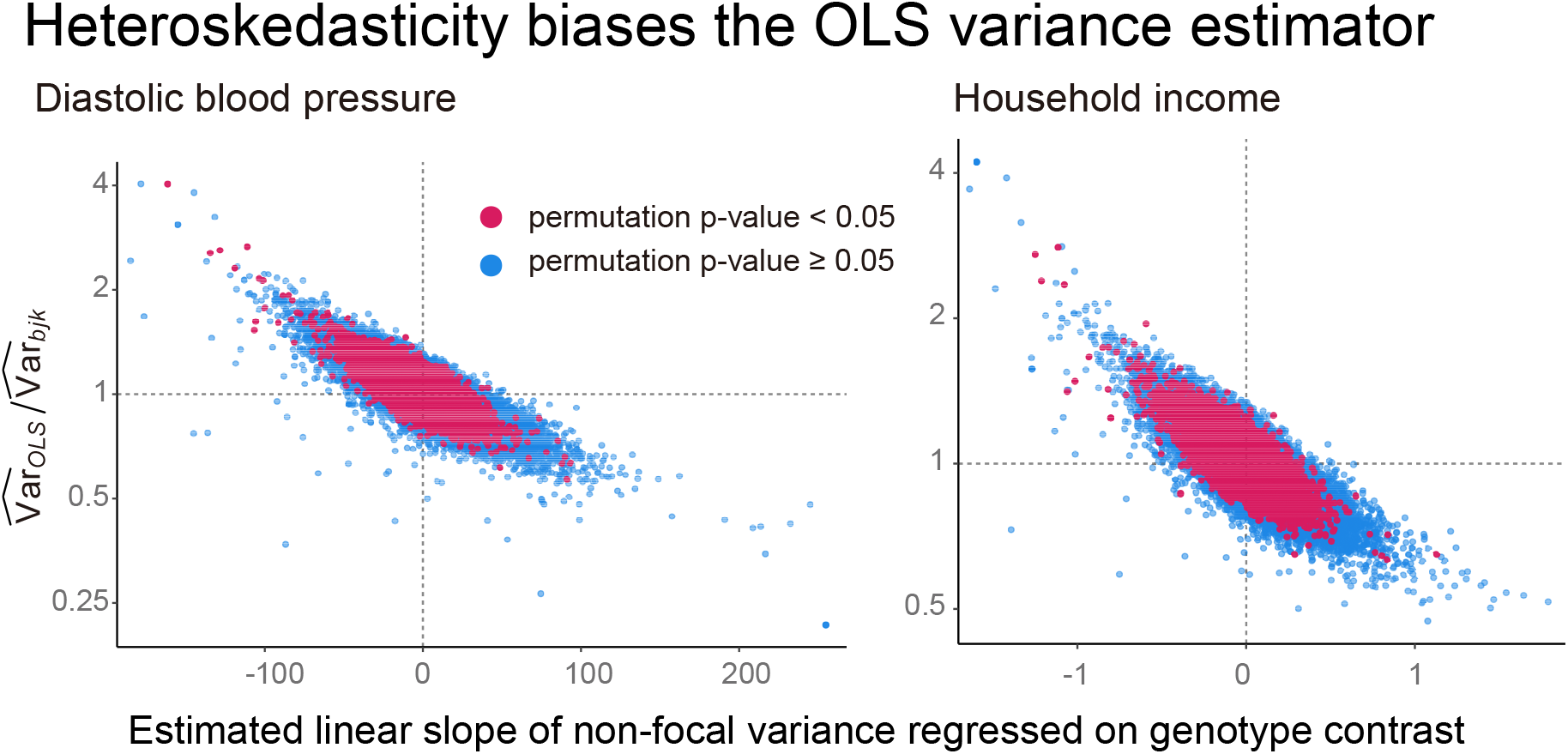
Heteroskedasticity factor drives the direction of the bias in the OLS variance estimator. To validate the analytical prediction in **Eq. 7**, that the OLS estimator is upward biased if the heteroskedasticity factor is negative and downward biased if the heteroskedasticity factor is positive, we compared the log_2_ ratio of OLS to block jackknife variance estimates against the estimated slope, which describes how non-focal variance changes with genotype contrast. The x-axis shows the estimated slope per SNP; the y-axis shows the log_2_ ratio of the OLS to block jackknife variance estimate, where values above and below 1 indicate upward and downward bias in the OLS estimator, respectively. Results are shown for diastolic blood pressure and household income as representative examples across the 11 physiological and 6 behavioral traits. Consistent with theoretical predictions, the ratios tended to be larger than 1 for negative slopes and smaller than 1 for positive slopes.

Overall, the empirical results from the UK Biobank closely mirrored our theoretical predictions. The permutation variance estimates were larger than the block jackknife benchmark for SNPs with evidence of non-zero allelic effects. The OLS variance estimator deviated from the block jackknife benchmark, driven by heteroskedastic non-focal variation. Both discrepancies were more pronounced for loci with smaller permutation *p*-values or rarer alleles.

## Discussion

Family-based studies provide estimates of direct genetic effects with minimal bias from population stratification or other confounding factors^1,9^. Although the biasing effects of stratification and assortative mating are largely removed by the family design^23^, the heterogeneity with which they are associated can still influence estimates of standard error.

We studied the behavior of three estimators of the (squared) standard error of allelic effect estimates in family GWAS. We focused on a setting with exactly two siblings per family in the main text, although we generalize this in the **Supplementary Material ??**. The first method, the usual OLS standard errors applied to a regression of within-sibship phenotypic differences on within-sibship genotypic differences, is biased in the presence of heteroskedasticity. We identify heterogeneity among families—including in genetic or environmental background— as an important driver of heteroskedasticity. The second method, using the standard deviation of the permutation distribution under the null, is biased when the null hypothesis is false, i.e., when the true allelic effect is nonzero. In contrast to these two estimators, block jackknife standard errors are approximately unbiased in the scenarios we studied and can be computed easily from the output of commonly-used bioinformatics tools (**Supplementary Material S5**). Finally, we confirmed our theoretical derivations with simulations and observed patterns consistent with them in biobank data.

We focus on the block jackknife here rather than leave-one-out jackknifing or bootstrapping for ease of computation. Standard leave-one-out jackknifing requires estimating the statistic of interest—here, the estimated allelic effect—*n* times, one for each of the *n* datasets that can be formed by leaving out one of the original observations. (In our setting, the observations are sibships, and *n* is the number of sibships.) In contrast, for block jackknife, we provided a guideline (**Eq. 10**) for choosing the number of block jackknife samples needed to achieve a given level of accuracy, managing a tradeoff between calibration and computational costs.

This reduced cost is important in GWAS contexts, where both the sample size and the number of genomic sites for which estimates are required can be very large.

The reason we focus on the jackknife over bootstrapping is more practical than computational *per se*: the jackknife is much easier to implement in *PLINK* ^29^, the most widely used software for GWAS. Whereas neither bootstrap nor jackknife standard errors for sibling-based GWAS are implemented in *PLINK*, it is easy to run *PLINK* analyses while excluding subjects in a user-provided list. Such analyses of subsets are the key ingredient of the jackknife and the basis for our analyses of UK Biobank data here. Were bootstrap methods designed for family data to be implemented directly in *PLINK* or other widely used software for GWAS, we expect that they would perform well. As shown in the **Supplementary Material S5**, the clustered standard errors used in some recent family studies^5^ also appear to be approximately unbiased, and they are computationally efficient, but they are not implemented in widely used software for family-GWAS, most notably *PLINK* ^29^ and *snipar* ^33^. In the meantime, resampling approaches like block jackknife are readily available.

We focused here on quantitative traits. However, we did not assume normality of errors, and we expect most of our results to carry over to binary traits. That said, when studying binary traits, often researchers prefer to estimate effects using a generalized linear model, such as logistic^37^ or probit regression^38^. Although our results do not translate directly to this setting, we expect block jackknife standard errors to perform well for GLM estimates as well. The issue perhaps requiring more serious investigation is ascertainment of families into a GWAS. Whereas we assume a population-based sample, studies of binary traits such as diseases often entail intensive sampling of cases^39,40^.

To understand the sources of genotype-phenotype associations, we will need to accurately quantify uncertainty in family GWAS, and we will need to be able to do this in workhorse software. Informed by our investigation of drivers of uncertainty in within-family GWAS, we argue that the block jackknife based measures of uncertainty are accurate, computationally efficient, and straightforward to compute with widely available tools.

## Acknowledgements

We thank Olivia Smith, Samuel Pattillo Smith, Liaoyi Xu, Charlotte LeMay, Thomas Juenger, Elliot Tucker-Drob, and members of the Harpak Lab for feedback on the manuscript. The work was funded by NIH grants R35GM137758 to M.D.E., and NIH grant R35GM151108 and a Pew Scholarship to A.H. This study was conducted using the UK Biobank resource under application 92741, as approved by the University of Texas at Austin institutional review board (study protocol 00003287). The authors acknowledge the Texas Advanced Computing Center (TACC) at The University of Texas at Austin for providing computational resources that have contributed to the research results reported within this paper.

## Methods

### The variance of the estimated allelic effect

In this section, we derive the variance of the estimated allelic effect in a sib-GWAS (**Eq. 5**). The estimator is the slope of an OLS regression of within-family phenotype differences (Δ *Y*) on within-family genotype differences (Δ *X*) across *n* families, following **Eq. 3**. The closed-form expression is:

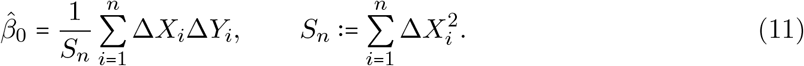

By the law of total variance:

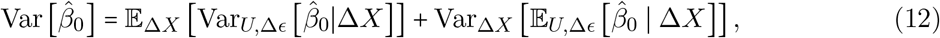

where Δ*X* =(Δ*X*_1_, …, Δ*X*_*n*_) is the vector of the within-family genotype differences at the focal allele; 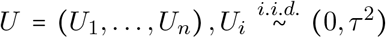 quantifies the heterogeneity of allelic effect *β*_*i*_(=*β*_0_ *U*_*i*_), and Δ*ϵ* (Δ*ϵ*_1_, …, Δ*ϵ*_*n*_) denotes family-specific differences in non-focal variation. We assume that the non-focal variation has mean zero and its variance changes linearly with the focal genotype contrast (**Eq. 4**). We allow a covariance between the allelic effect and non-focal variation: Cov [*U*_*i*_, Δ *ϵ* _*i*_ | Δ *X*_*i*_] =*c*_*i*_ and show below that this covariance does not contribute to the variance of the allelic effect estimates. We assume independence among families,

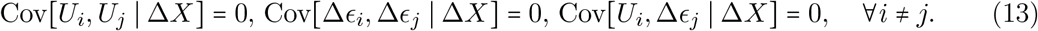

We derive the two terms of **Eq. 12** in turn. To derive the first term, we expand **Eq. 11** by substituting **Eq. 3**,

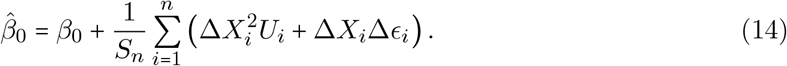

Substituting **Eq. 14** into the inner conditional variance of the first term in **Eq. 12** yields

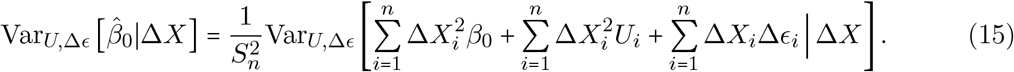

The first term, 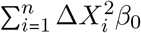, is constant given the genotype difference Δ *X*, and thus contributes no variance. Hence, **Eq. 15** reduces to:

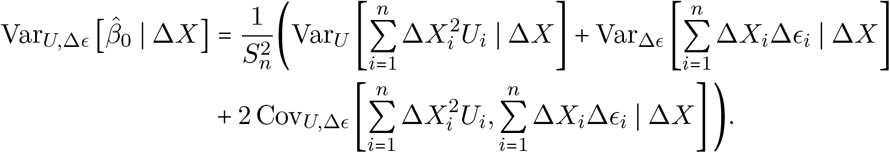

By independence across families (**Eq. 13**), all cross-family covariances vanish:

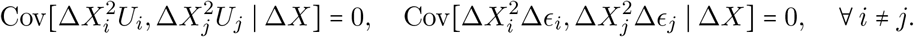

Additionally, using

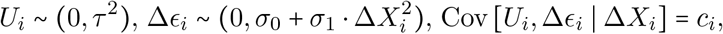

**Eq. 15** is:

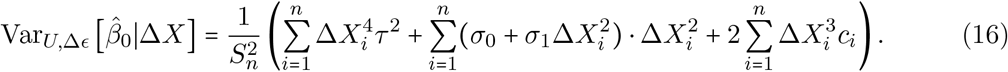

To further simplify **Eq. 16**, we define the leverage 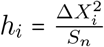, which reflects the influence of the *i*-th sibling pair on the regression line: pairs with larger genotype difference (Δ *X*_*i*_) exert higher leverage on 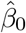. **Eq. 16** can then be written as:

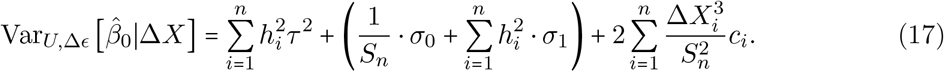

Next, we take the outer expectation with respect to genotype difference Δ *X* in **Eq. 17**. Assuming Hardy-Weinberg equilibrium, the following expectations hold (a derivation is given in the **Supplementary Material S1**):

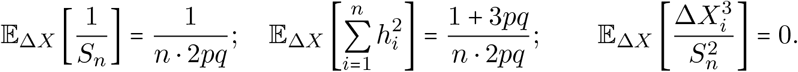

Taking the outer expectation with respect to genotype difference then gives the first term of the total variance (**Eq. 12**):

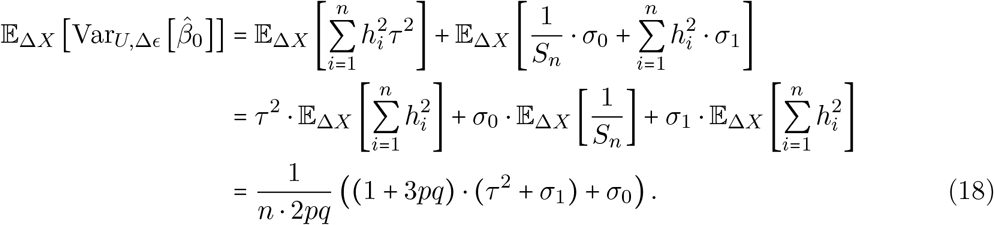

We next focus on the second term of the total variance (**Eq. 12**). Since both *U*_*i*_ and Δ *ϵ*_*i*_ have mean zero, the conditional expectation of the second term in **Eq. 12** is:

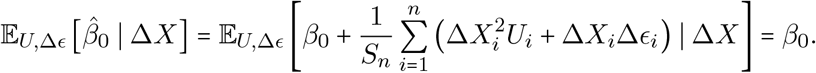

Since *β*_0_ is a constant, the outer variance is zero:

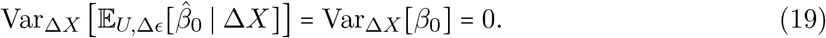

Finally, substituting **Eq. 18** and **Eq. 19** into the total variance (**Eq. 12**) yields the variance of the allelic effect estimator:

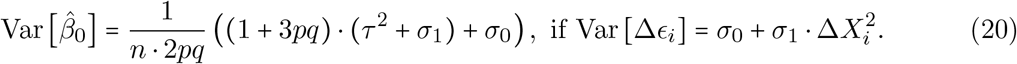

### Derivation of the bias of the OLS variance estimator

In this section, under the assumption that the non-focal variance changes linearly with the genotype contrast (**Eq. 4**), we derive the bias of the OLS variance estimator:

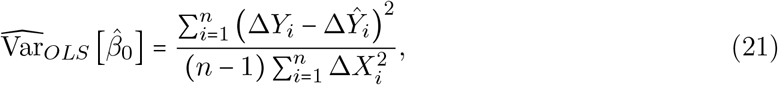

where Δ *Y*_*i*_ is the observed phenotype difference in family *i* and 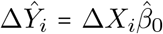 is the predicted value. We derive the expectation of the OLS variance estimator (**Eq. 21**) and compare it with the true variance of 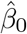 (**Eq. 5**) to obtain its bias.

First, we expand the OLS variance estimator into variables with known distributions. Substituting the additive model **Eq. 3** and the explicit form of 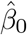 (**Eq. 14**) decomposes Δ *Y*_*i*_ − Δ *Ŷ* _*i*_ into two terms:

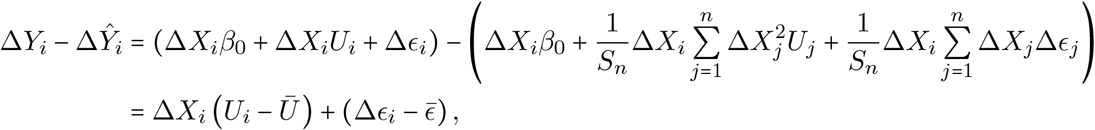

with 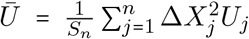 and 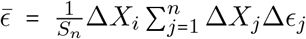. Squaring and summing yields the residual sum of squares:

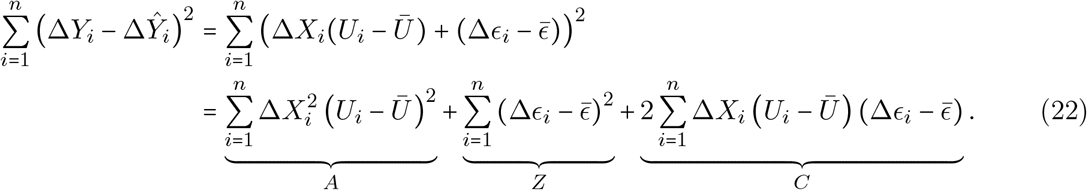

The three terms in **Eq. 22** can be simplified as:

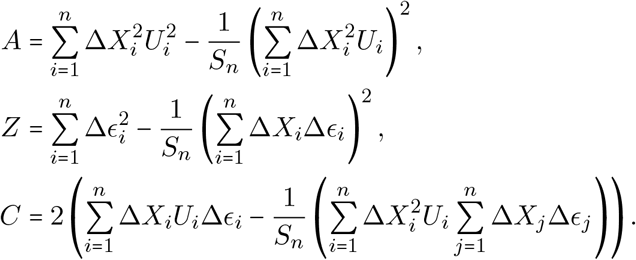

Substituting these simplified forms into **Eq. 21** expands the OLS variance estimator into:

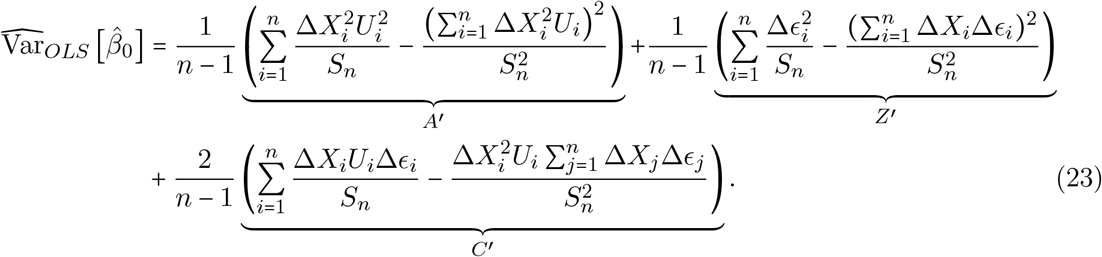

After expanding the OLS variance estimator (**Eq. 21**) into variables with known distributions (**Eq. 23**), we derive its expectation using the Law of Total Expectation,

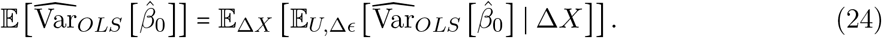

We evaluate this expectation (**Eq. 24**) term by term. Since 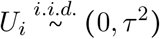, we have

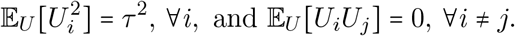

The conditional expectation of *A*^′^ is:

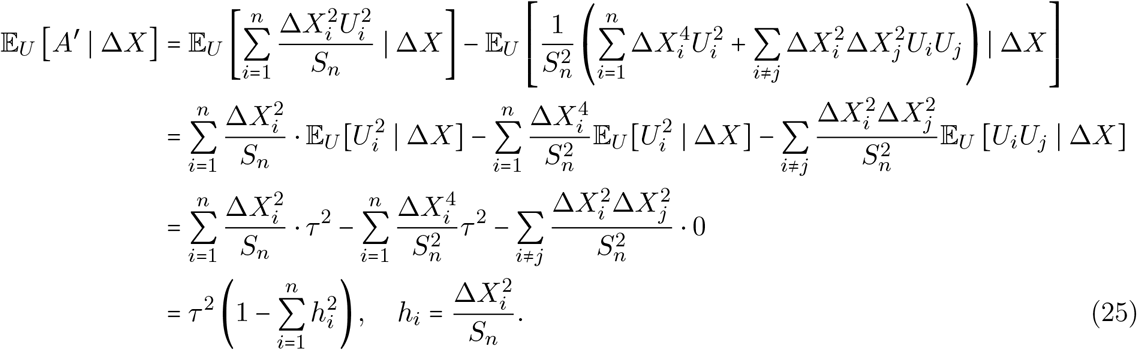

In the **Supplementary Material S1**, we show that 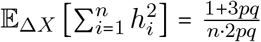. Therefore, taking expectation with respect to Δ*X* in **Eq. 25** gives the total expectation of *A*^′^:

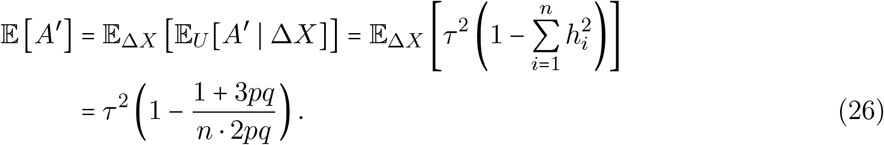

We next derive the expectation of term *Z*^′^ in **Eq. 23**. Recall from **Eq. 4** that

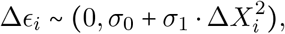

and that Δ*ϵ*_*i*_ are mutually independent across families. It follows that

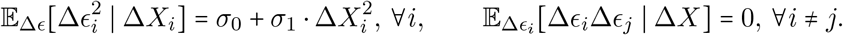

Therefore, the conditional expectation of the *Z*^′^ term is:

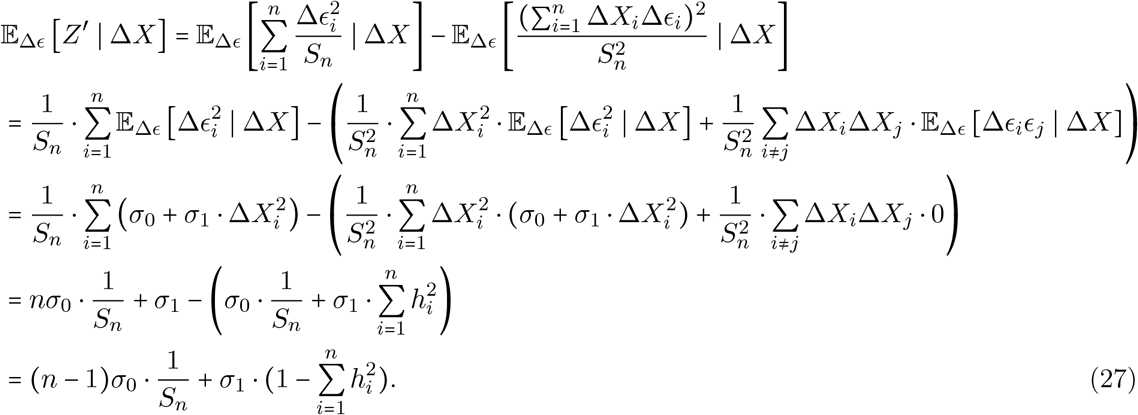

We show in the **Supplementary Material S1** that:

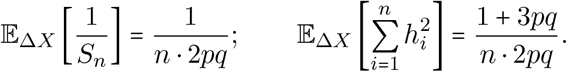

Then, taking the outer expectation with respect to Δ *X* in **Eq. 27** gives the total expectation of *Z*^′^,

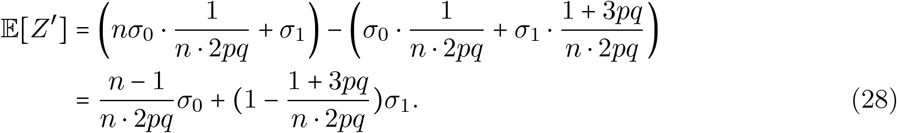

We next show that the expectation of the term in **Eq. 23** is zero. The within-family covariance Cov [*β*_*i*_, Δ ϵ _*i*_] = *c*_*i*_ is permitted, while cross-family covariances are assumed to be zero (**Eq. 13**). The conditional expectation of *C*^′^ is therefore:

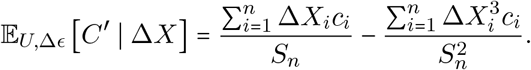

Since 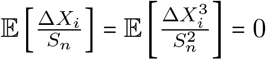, for all *i* (**Supplementary Material S1**),

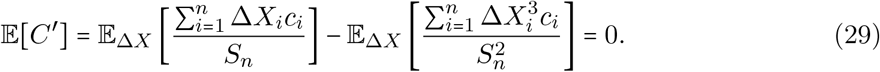

Combining **Eqs. 26, 28, 29** and substituting into **Eq. 23**,

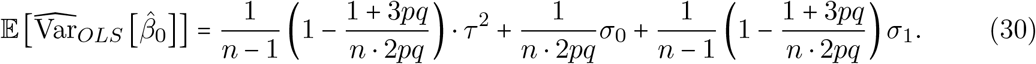

**Eq. 30** shows that the expectation of the OLS variance estimator is determined by both the heterogeneity of the allelic effect (**Eq. 2**) and the dependence between non-focal variance and focal genotype contrast (**Eq. 4**). Comparing **Eq. 30** and the true variance of 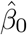 (**Eq. 18**) yields the bias of the OLS variance estimator:

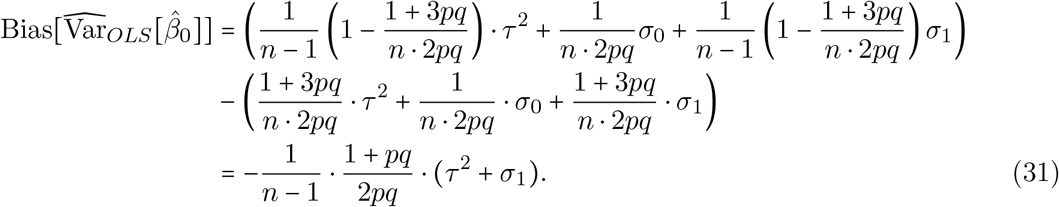

### Derivation of the bias of the permutation variance estimator

In this section, we derive the bias of the permutation variance estimator. Unlike the OLS case, which required a specific functional form for the heteroskedasticity (**Eq. 4**), the permutation bias derivation holds under the general assumption that the non-focal variance depends arbitrarily on the focal genotype contrast:

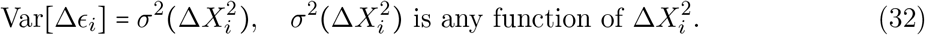

The bias expression for the permutation estimator (**Eq. 8**) holds regardless of how the non-focal variance scales with genotype contrast.

The permutation variance estimator uses the sample variance of the allelic effect estimates across permutation replicates. In the sib-GWAS with two siblings per family, the permutation is equivalent to randomly resampling the sign of the within-family phenotype differences. Specifically, for each family *i* and permutation replicate *b*, we multiply Δ *Y*_*i*_ by an independent sign variable 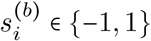, with 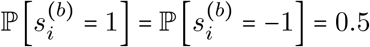 and 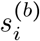 drawn independently across families. The allelic effect estimator in permutation replicate *b* is then:

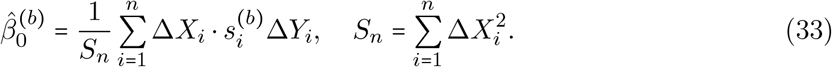

The sample variance across replicates gives the permutation variance estimator:

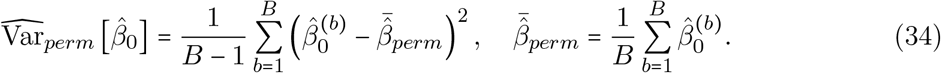

By the Law of Large Numbers (LLN), as the number of permutation replicates increases to infinity, *B* → ∞, the sample variance converges to the population variance of 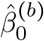:

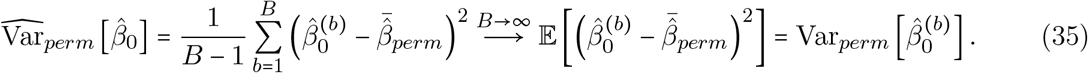

By the law of total variance, the right-hand side of **Eq. 35** is:

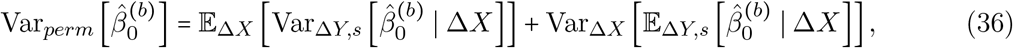

where 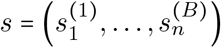 denotes the full collection of sign variables across replicates and families.

We start by proving that the second term in **Eq. 36** vanishes. Since 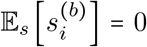 and 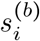 is independent of the sibling information, 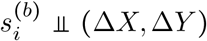, the inner conditional expectation satisfies:

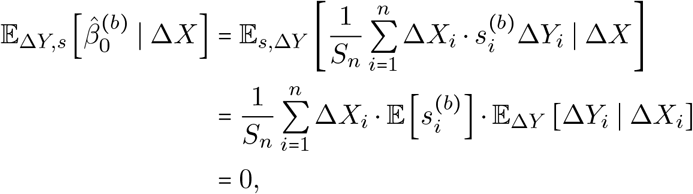

where the product vanishes because 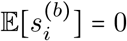 regardless of the value of E [Δ *Y*_*i*_ ∣ Δ *X*_*i*_]. Therefore, **Eq. 36** reduces to:

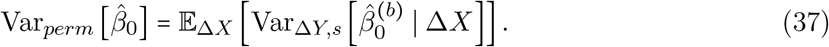

Substituting **Eq. 33** into the inner conditional variance of **Eq. 37**:

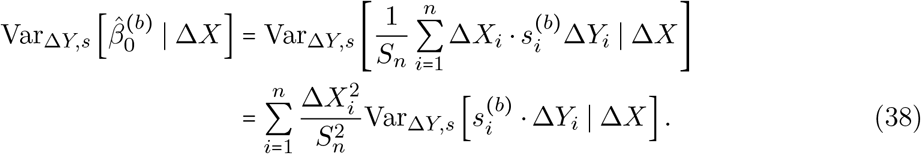

Since 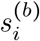 is independent of the sibling phenotypes 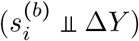 and 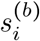 has mean 0 and variance 1, the conditional variance in **Eq. 38** is:

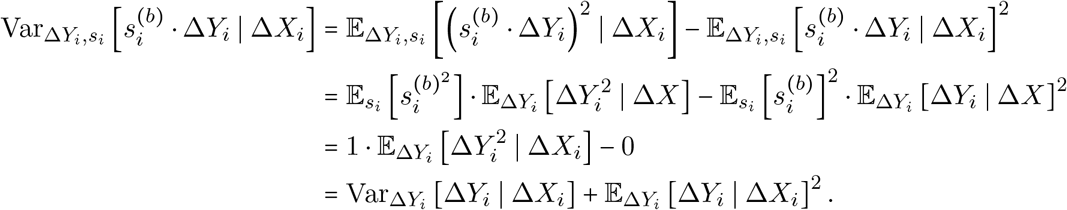

Using the additive model (**Eq. 3**) and 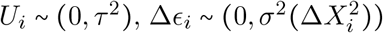:

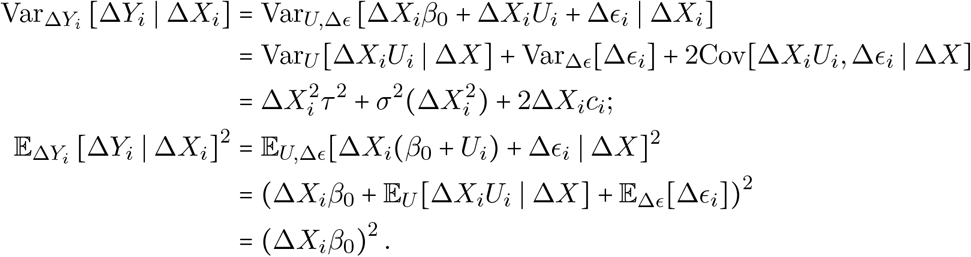

Therefore, the inner conditional variance of **Eq. 37** is:

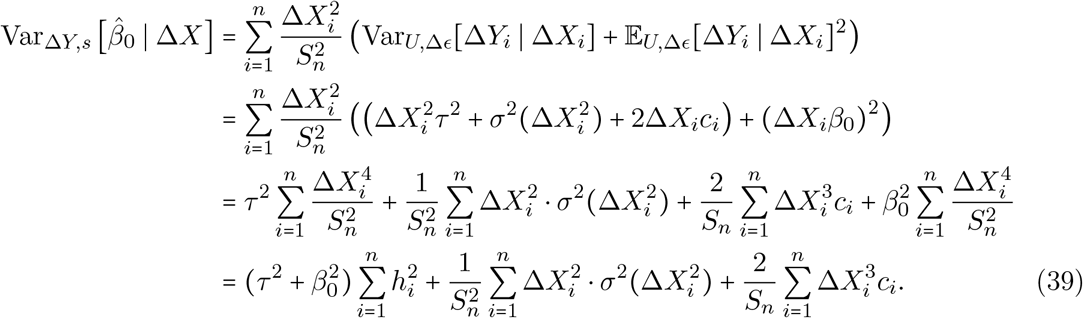

We show in the **Supplementary Material S1** that under Hardy-Weinberg Equilibrium,

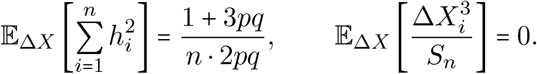

Taking the expectation with respect to Δ *X* in **Eq. 39**,

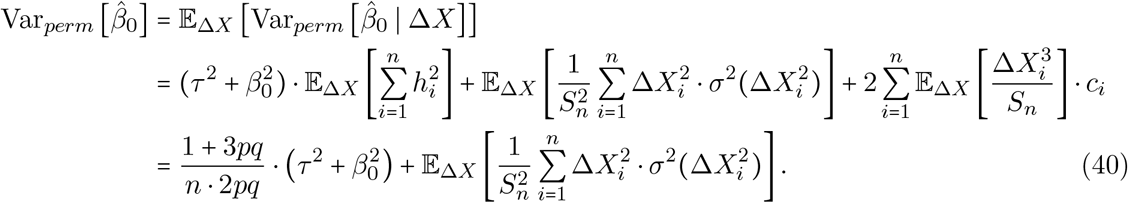

To compare with **Eq. 40**, we derive the true variance Var 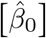 under the general heteroskedasticity assumption (**Eq. 32**). Following the same argument as in the true-variance derivation but replacing **Eq. 4** with **Eq. 32** in **Eq. 15** gives

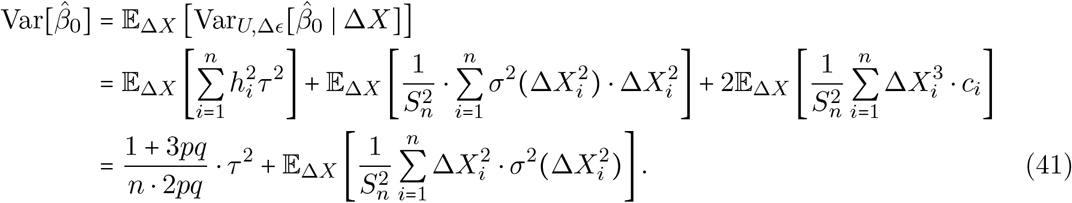

Subtracting **Eq. 41** from **Eq. 40** gives the bias of the permutation estimator:

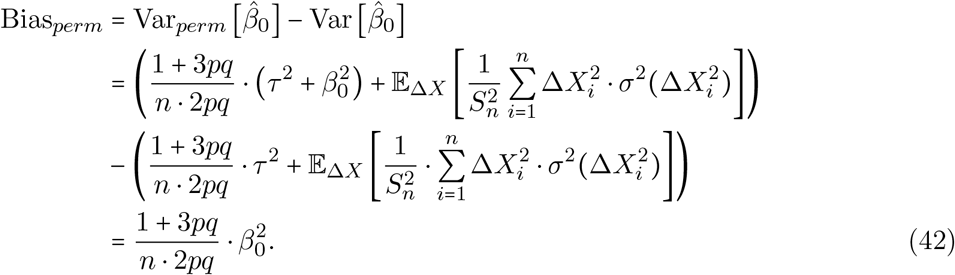

The two heteroskedasticity-dependent terms 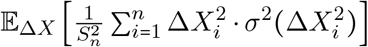 cancel exactly, leaving a bias that depends only on the baseline allelic effect *β*_0_.

### Derivation of the bias of the block jackknife variance estimator

In this section, we derive the finite-sample bias of the block jackknife variance estimator. In each block jackknife resample, a block of *d* sibling pairs is randomly omitted from the full sample of size *n*, leaving *r* = *n* − *d* pairs. For each of *M* independent replicates, let *S*_*m*_ ⊂ {1, …, *n*} denote the retained indices with cardinality *r*. The corresponding re-estimated allelic effect is:

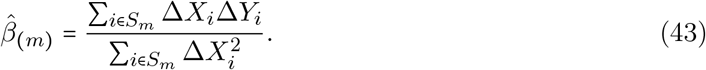

After *M* replicates, we estimate the variance of 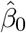 using the sample variance of the *M* re-estimated allelic effects:

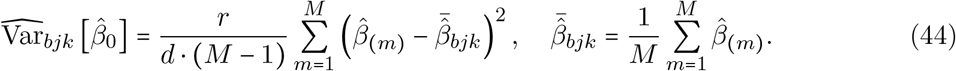

To derive its expectation, we first express 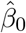 as a smooth function of a sample mean and apply a Taylor expansion, following Section 2.4 of Shao and Tu^41^. We define the observation vector 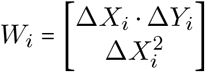, with the mean of 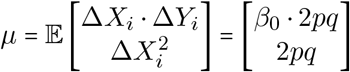. Then, the allelic effect estimator 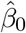 can be written as a smooth function of the sample mean:

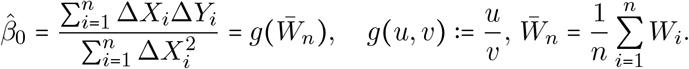

Expanding 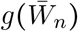 around *µ*:

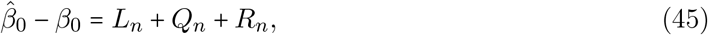

where *L*_*n*_ is the first-order term, *Q*_*n*_ is the second-order term, and the remainder *R*_*n*_ = *o*_*p*_(*n*^−1^). Specifically,

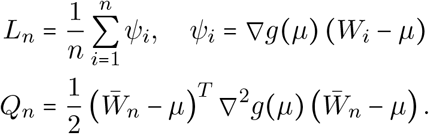

Under standard moment conditions, *L*_*n*_ = *O*_*p*_ (*n*^−1/ 2^) and *Q*_*n*_ = *O*_*p*_ (*n*^−1^). Therefore, following **Eq. 45**, the variance of 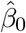 is:

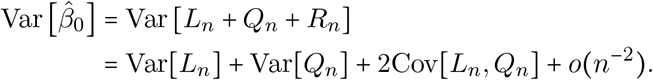

We show in the **Supplementary Material S2.1** that the second-order term Var (*Q*_*n*_) = *O* (*n*^−2^) and the covariance term Cov [*L*_*n*_, *Q*_*n*_] =(*O n*^−3^/^2^). The leading contribution comes from the first-order term *L*_*n*_. Hence,

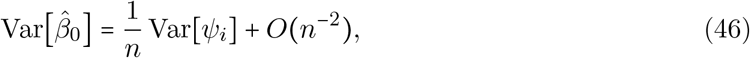

Having derived 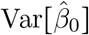 using the Taylor expansion, we next apply the same argument to each block jackknife replicate. For the (*m*)-th resample, the re-estimated allelic effect decomposes as:

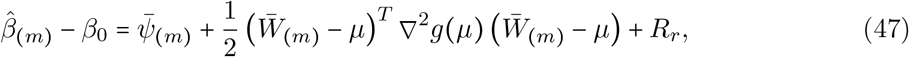

where

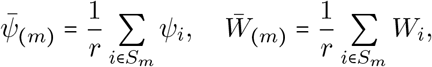

and *R*_*r*_ = *o* _*p*_ (*r*^−1^). In each block jackknife resample, each sibling pair is retained with probability 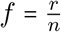. Hence,

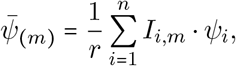

where *I*_*i,m*_ is an indicator variable equal to 1 if the *i*-th sibling pair is retained in block jackknife resample *m* and 0 otherwise. Under simple random sampling without replacement, the indicator variables satisfy:

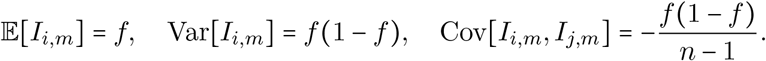

By the law of total expectation:

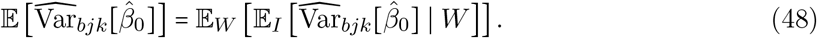

Since the sample variance across replicates is an unbiased estimator of the true conditional variance 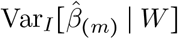,

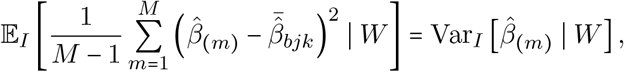

the inner conditional expectation in **Eq. 48** simplifies to:

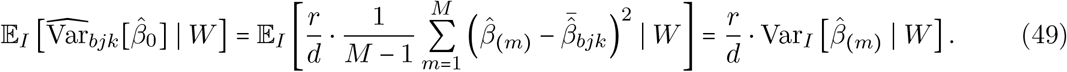

Substituting the Taylor expansion (**Eq. 47**) into the conditional variance (**Eq. 49**) gives the conditional variance with respect to random sampling:

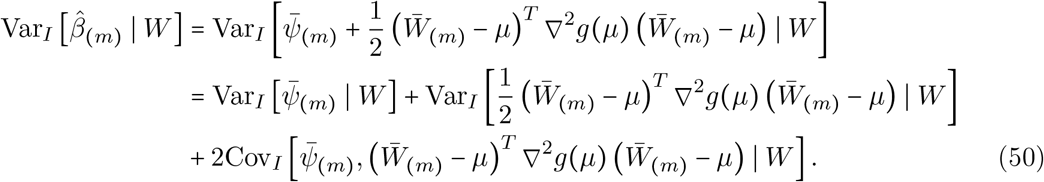

The first term provides the leading contribution. Using the indicator moments:

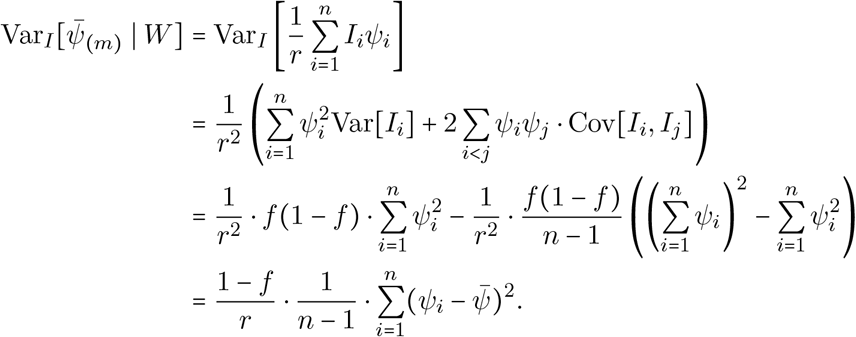

In the **Supplementary Material S2.2**, we show that the second-order variance term in Eq. 50 is of order *O*(*r*^−2^) and the covariance term is of order *O*(*r*^−3/2^); both are negligible relative to the *O*(*r*^−1^) leading term. Substituting into Eq. 49,

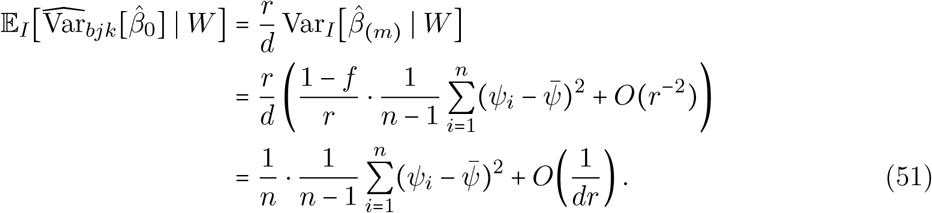

Taking the expectation with respect to *W* in Eq. 51, we obtain the expectation of the block jackknife variance estimator:

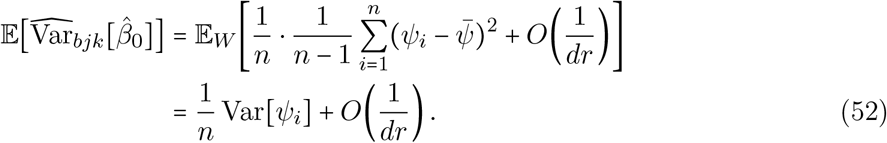

The expectation of the block jackknife variance estimator matches the leading *n*^−1^ term in 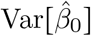, up to a higher-order discrepancy. Subtracting **Eq. 46** from **Eq. 52** yields the bias of the block jackknife variance estimator:

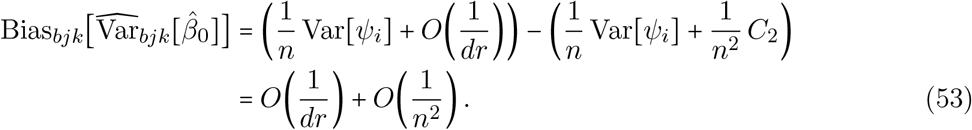

Both terms vanish as *n* grows, establishing that the block jackknife estimator is asymptotically unbiased.

### Decision rule for the number of replicates required by the block jackknife method

The block jackknife provides an asymptotically unbiased estimate of 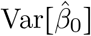, while avoiding the computational burden of full jackknife resampling. To determine how many resampling replicates are needed for a stable estimate, we derive a lower bound that ensures the block jackknife variance estimate is close to the true value with high probability.

Suppose we perform *m* independent resampling replicates. The block jackknife variance estimator is:

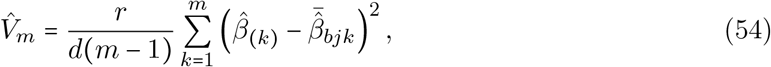

where 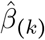 is the allelic effect estimated in the *k*-th replicate, and 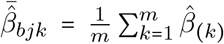 is the average estimated effect across all replicates.

Let 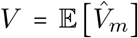 denote the expectation of the block jackknife variance estimator over the resampling distribution. As shown in the previous section, *V* approximates the true variance of 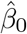.

By Chebyshev’s inequality, the probability that the block jackknife variance estimator 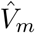 deviates from its expectation *V* by more than *ϵ* > 0 satisfies:

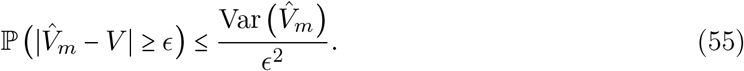

Setting *ϵ* = *δ V* for a relative tolerance *δ* > 0 gives:

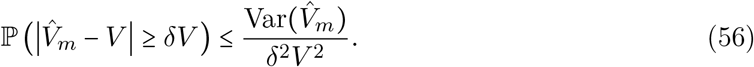

To evaluate the right-hand side, we treat the replicate estimates 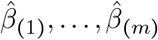, as approximately independent and identically distributed draws from the resampling distribution. Under this approximation, the variance of the sample variance 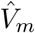 is related to the kurtosis of 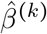 via:

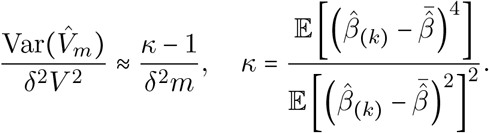

By the central limit theorem, 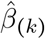 is approximately normal for large *r*, giving *κ* ≈ 3 and thus *κ* −1 ≈ 2. Substituting the approximation into **Ineq. 56**, we obtain:

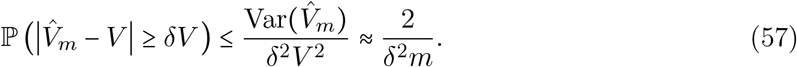

To translate this bound from the variance estimate 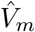 to the estimated standard error 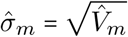, we use 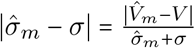. Since 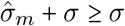:

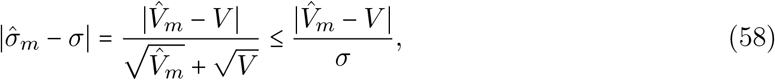

and therefore:

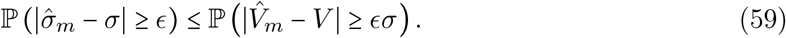

Combining **Eq. 57** and **Eq. 59** with *ϵ* = *δ* ⋅ *σ*, the probability that the block jackknife standard error estimate deviates from the true value by more than a *δ*−fraction is bounded by 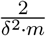. To ensure this probability does not exceed *α*, it suffices to choose:

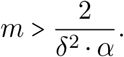

For instance, to guarantee that the estimated standard error is within *δ* = 20% of the true value with at least 1 − *α* = 90%, it requires 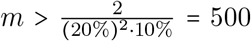 replicates. Our empirical analyses used *m* = 500, which satisfies this criterion.

### Application to UK Biobank data

We applied the three variance estimators to sib-GWAS data from the UK Biobank^35^. The cohort construction, phenotype preprocessing, and genotype quality control followed the pipeline described in Smith, Smith et al.^9^ and are summarized here. Sibling pairs were identified from the relatedness information provided by the UK Biobank: pairs were retained if their kinship coefficient estimate fell between 0.1768 and 0.3536, and their IBS0 value exceeded 0.0012, as estimated by *KING* ^42^. From each resulting family group, two siblings were selected, yielding 17,353 White British full sibling pairs in which each individual belongs to exactly one pair. We analyzed 17 continuous phenotypes, spanning 11 physiological traits and 6 behavioral traits. Following Smith, Smith et al.^9^, we residualized individuals’ phenotypes for age and sex. Sample sizes per trait ranged from 2,163 to 17,328 sibling pairs after removing pairs with missing data.

Genotypic variants were filtered using *plink2* ^43^. Specifically, variants with INFO score below 0.8, genotype missingness above 0.05 (`–geno 0.05`), minor allele count of 5 or fewer (`–mac 5`), non-biallelic sites), or Hardy-Weinberg equilibrium *p* <10^−10^(`–hwe 1e-10`) were excluded, leaving 9,607,691 variants. We focused on a set of SNPs obtained by clumping on linkage disequilibrium (*r*^2^ < 0.1, window = 100kb), without thresholding on *p*-value. This procedure ensures that the analyzed SNPs span a broad, approximately unselected range of allelic effect sizes, allowing assessment of estimator behavior from null to strongly associated SNPs. Clumping was performed within the sibling cohort, leaving 504,858 SNPs for analysis.

Each of the three variance estimators was then computed for every SNP-trait combination, using the procedure described below. To obtain the OLS variance estimates, we performed OLS regression of within-family phenotype differences on within-family genotype differences, following the additive model (**Eq. 3**). The variance, 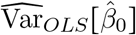, was computed analytically from the RSS of this regression, using custom scripts in *R*^44^.

To obtain the permutation variance estimates, we used the standard permutation procedure in *PLINK1*.*9*, performing 1,000 replicates per SNP in which phenotype labels were randomly swapped within sibships. The variance of the resulting empirical distribution of allelic effect estimates serves as the permutation-based variance estimate. To rank SNPs by strength of evidence for a nonzero allelic effect in **Figs. 3 (A-B), S7, S8**, we obtained *p*-values via adaptive permutation in *PLINK1*.*9* with a minimum of 1,000 and a maximum of 1,000,000 permutations per SNP.

For the block jackknife variance estimates, we conducted 500 resampling replicates per SNP, each omitting *d* = 500 sibling pairs sampled at random without replacement. The block jackknife variance estimate was then computed from the variance of the allelic effect estimates across these 500 subsamples, following **Eq. 44**.

To quantify the discrepancy between the OLS (and permutation estimators) and the block jackknife benchmark, we computed the log_2_ ratio of each estimator to the block jackknife estimate for each SNP, pooled across all 17 traits. SNPs were binned by the estimated *p*-value of the allelic effect (**Figs. 3(A-B), S7, S8**) or by minor allele frequency (**Figs. 3(C-D), S9, S10**), with each bin containing 300 SNPs.

### Estimating the slope describing how non-focal variance changes with focal genotype contrast

To test the prediction of **Eq. 7** empirically, we constructed a per-SNP estimate of *σ*_1_, the slope in **Eq. 4** describing how non-focal variance changes with focal genotype contrast.

For each SNP, we partitioned the *n* sibling pairs into 3 groups according to their focal genotype contrast 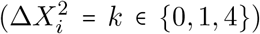, which are the only values 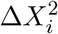 can take under biallelic Mendelian segregation. Within each group, we estimated the non-focal variance using the regression residuals 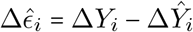:

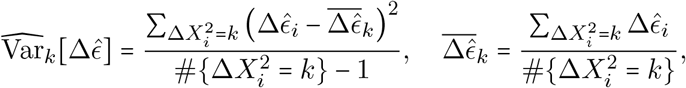

We then regressed the three group-level non-focal variance estimates on their corresponding genotype contrast values to obtain 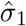:

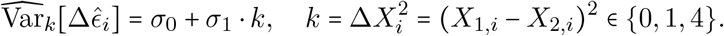

## Code availability

All summary statistics generated in this study are available on the Harpak Lab website’s data tab https://www.harpaklab.com/data. The code used to generate all results and figures is available at https://github.com/harpak-lab/Measure-uncertainty-in-sib-GWAS.

## References

[1] Alexander I. Young, Stefania Benonisdottir, Molly Przeworski, and Augustine Kong. “Deconstructing the sources of genotype-phenotype associations in humans”. Science 365.6460 (2019), pp. 1396–1400. ISSN: 1095-9203. DOI: 10.1126/science.aax3710.

[2] Jurg Ott, Jing Wang, and Suzanne M. Leal. “Genetic linkage analysis in the age of whole-genome sequencing”. Nature Reviews Genetics 16.5 (Mar. 2015), pp. 275–284. ISSN: 1471-0064. DOI: 10.1038/nrg3908.

[3] Warren J Ewens and Richard S Spielman. “The transmission/disequilibrium test: history, subdivision, and admixture”. American journal of human genetics 57.2 (1995), p. 455.

[4] Neil Risch and Kathleen Merikangas. “The Future of Genetic Studies of Complex Human Diseases”. Science 273.5281 (1996), pp. 1516–1517. ISSN: 1095-9203. DOI: 10.1126/science.273.5281.1516.

[5] Laurence J. Howe et al. “Within-sibship genome-wide association analyses decrease bias in estimates of direct genetic effects”. Nature Genetics 54.5 (May 2022), pp. 581–592. ISSN: 1546-1718. DOI: 10.1038/s41588-022-01062-7.

[6] Tammy Tan et al. “Family-GWAS reveals effects of environment and mating on genetic associations” (Oct. 2024). DOI: 10.1101/2024.10.01.24314703.

[7] Matthew R Robinson et al. “Population genetic differentiation of height and body mass index across Europe”. Nature Genetics 47.11 (2015), pp. 1357–1362. ISSN: 1546-1718. DOI: 10.1038/ng.3401.

[8] Hakhamanesh Mostafavi, Arbel Harpak, Ipsita Agarwal, Dalton Conley, Jonathan K Pritchard, and Molly Przeworski. “Variable prediction accuracy of polygenic scores within an ancestry group”. elife 9 (2020), e48376.

[9] Samuel Pattillo Smith et al. “A Litmus Test for Confounding in Polygenic Scores” (Feb. 2025). DOI: 10.1101/2025.02.01.635985.

[10] Jeremy J Berg et al. “Reduced signal for polygenic adaptation of height in UK Biobank”. eLife 8 (Mar. 2019). ISSN: 2050-084X. DOI: 10.7554/eLife.39725.

[11] Mashaal Sohail et al. “Polygenic adaptation on height is overestimated due to uncorrected stratification in genome-wide association studies”. eLife 8 (Mar. 2019). ISSN: 2050-084X. DOI: 10.7554/eLife.39702.

[12] Nick Barton, Joachim Hermisson, and Magnus Nordborg. “Why structure matters”. eLife 8 (Mar. 2019). ISSN: 2050-084X. DOI: 10.7554/eLife.45380.

[13] Augustine Kong et al. “The nature of nurture: Effects of parental genotypes”. Science 359.6374 (Jan. 2018), pp. 424–428. ISSN: 1095-9203. DOI: 10.1126/science.aan6877.

[14] Michel G. Nivard et al. “More than nature and nurture, indirect genetic effects on children’s academic achievement are consequences of dynastic social processes”. Nature Human Behaviour 8.4 (Jan. 2024), pp. 771–778. ISSN: 2397-3374. DOI: 10.1038/s41562-023-01796-2.

[15] Arbel Harpak and Michael D. Edge. “GWAS deems parents guilty by association”. Proceedings of the National Academy of Sciences 118.27 (2021). ISSN: 1091-6490. DOI: 10.1073/pnas.2109433118.

[16] Per Magnus et al. “Cohort Profile Update: The Norwegian Mother and Child Cohort Study (MoBa)”. International Journal of Epidemiology 45.2 (Apr. 2016), pp. 382–388. ISSN: 1464-3685. DOI: 10.1093/ije/dyw029.

[17] Yuchang Wu et al. “Estimating genetic nurture with summary statistics of multigenerational genome-wide association studies”. Proceedings of the National Academy of Sciences 118.25 (2021). ISSN: 1091-6490. DOI: 10.1073/pnas.2023184118.

[18] Nicole M Warrington et al. “Maternal and fetal genetic effects on birth weight and their relevance to cardio-metabolic risk factors”. Nature genetics 51.5 (2019), pp. 804–814.

[19] Loic Yengo et al. “Imprint of assortative mating on the human genome”. Nature Human Behaviour 2.12 (Nov. 2018), pp. 948–954. ISSN: 2397-3374. DOI: 10.1038/s41562-018-0476-3.

[20] Alexander Strudwick Young. “Estimation of indirect genetic effects and heritability under assortative mating” (2023). DOI: 10.1101/2023.07.10.548458.

[21] Yair Field et al. “Detection of human adaptation during the past 2000 years”. Science 354.6313 (Nov. 2016), pp. 760–764. ISSN: 1095-9203. DOI: 10.1126/science.aag0776.

[22] Michael D Edge and Graham Coop. “Reconstructing the History of Polygenic Scores Using Coalescent Trees”. Genetics 211.1 (Nov. 2018), pp. 235–262. ISSN: 1943-2631. DOI: 10.1534/genetics.118.301687.

[23] Carl Veller and Graham M. Coop. “Interpreting population- and family-based genome-wide association studies in the presence of confounding”. PLOS Biology 22.4 (Apr. 2024). Ed. by Priya Moorjani, e3002511. ISSN: 1545-7885. DOI: 10.1371/journal.pbio.3002511.

[24] Iftikhar J. Kullo, Cathryn M. Lewis, Michael Inouye, Alicia R. Martin, Samuli Ripatti, and Nilanjan Chatterjee. “Polygenic scores in biomedical research”. Nature Reviews Genetics 23.9 (Mar. 2022), pp. 524–532. ISSN: 1471-0064. DOI: 10.1038/s41576-022-00470-z.

[25] Jian Yang, S. Hong Lee, Michael E. Goddard, and Peter M. Visscher. “GCTA: A Tool for Genome-wide Complex Trait Analysis”. The American Journal of Human Genetics 88.1 (Jan. 2011), pp. 76–82. ISSN: 0002-9297. DOI: 10.1016/j.ajhg.2010.11.011.

[26] Brendan K. Bulik-Sullivan et al. “LD Score regression distinguishes confounding from polygenicity in genome-wide association studies”. Nature Genetics 47.3 (Feb. 2015), pp. 291–295. ISSN: 1546-1718. DOI: 10.1038/ng.3211.

[27] Brendan Bulik-Sullivan et al. “An atlas of genetic correlations across human diseases and traits”. Nature Genetics 47.11 (2015), pp. 1236–1241. ISSN: 1546-1718. DOI: 10.1038/ng.3406.

[28] Ben Brumpton et al. “Avoiding dynastic, assortative mating, and population stratification biases in Mendelian randomization through within-family analyses”. Nature Communications 11.1 (2020). ISSN: 2041-1723. DOI: 10.1038/s41467-020-17117-4.

[29] Shaun Purcell et al. “PLINK: A Tool Set for Whole-Genome Association and Population-Based Linkage Analyses”. The American Journal of Human Genetics 81.3 (2007), pp. 559–575. ISSN: 0002-9297. DOI: 10.1086/519795.

[30] Carl Veller, Molly Przeworski, and Graham Coop. “Causal interpretations of family GWAS in the presence of heterogeneous effects”. Proceedings of the National Academy of Sciences 121.38 (2024). ISSN: 1091-6490. DOI: 10.1073/pnas.2401379121.

[31] Jun Shao and CF Jeff Wu. “A General Theory for Jackknife Variance Estimation”. The Annals of Statistics 17.3 (1989), pp. 1176–1197.

[32] Bradley Efron and Charles Stein. “The Jackknife Estimate of Variance”. The Annals of Statistics 9.3 (May 1981). ISSN: 0090-5364. DOI: 10.1214/aos/1176345462.

[33] Alexander I. Young et al. “Mendelian imputation of parental genotypes improves estimates of direct genetic effects”. Nature Genetics 54.6 (2022), pp. 897–905. ISSN: 1546-1718. DOI: 10.1038/s41588-022-01085-0.

[34] Bradley Efron and Robert J Tibshirani. An introduction to the bootstrap. Chapman and Hall/CRC, 1994.

[35] Clare Bycroft et al. “The UK Biobank resource with deep phenotyping and genomic data”. Nature 562.7726 (Oct. 2018), pp. 203–209. ISSN: 1476-4687. DOI: 10.1038/s41586-018-0579-z.

[36] Joyce Y Wang et al. “Three open questions in polygenic score portability”. Nature Communications 17.1 (2026), p. 942.

[37] David R Cox. “The regression analysis of binary sequences”. Journal of the Royal Statistical Society Series B: Statistical Methodology 20.2 (1958), pp. 215–232.

[38] Chester I Bliss. “The method of probits”. Science 79.2037 (1934), pp. 38–39.

[39] Sanjana M Paye and Michael D Edge. “Mathematical bounds on r2 and the effect size in case-control genome-wide association studies”. Theoretical Population Biology 164 (2025), pp. 1–11.

[40] Jared M. Cole, Shane Rybacki, Samuel Pattillio Smith, Olivia S. Smith, and Arbel Harpak. “Representation in genetic studies affects inference about genetic architecture” (Jan. 2026). DOI: 10.64898/2026.01.12.699135.

[41] Jun Shao and Dongsheng Tu. “Theory for the Jackknife”. In: The Jackknife and Bootstrap. Springer New York, 1995, pp. 23–70. ISBN: 9781461207955. DOI: 10.1007/978-1-4612-0795-5_2.

[42] Ani Manichaikul, Josyf C Mychaleckyj, Stephen S Rich, Kathy Daly, Michèle Sale, and Wei-Min Chen. “Robust relationship inference in genome-wide association studies”. Bioinformatics 26.22 (2010), pp. 2867–2873.

[43] Christopher C Chang, Carson C Chow, Laurent CAM Tellier, Shashaank Vattikuti, Shaun M Purcell, and James J Lee. “Second-generation PLINK: rising to the challenge of larger and richer datasets”. Gigascience 4.1 (2015), s13742–015.

[44] Ross Ihaka and Robert Gentleman. “R: a language for data analysis and graphics”. Journal of computational and graphical statistics 5.3 (1996), pp. 299–314.

