## Supplementary Materials for "Estimating uncertainty in family-based GWAS"

August 21, 2026

### S1 Derivation of the expectation of genotype difference

In the main text, we used the following expectations to derive both the variance of the allelic effect estimator (**Eq. 5**) and the biases of variance estimators (**Eq. 21, Eq. 34**):

$$\mathbb{E}_{\Delta X}\left[\frac{1}{S_n}\right] = \frac{1}{n \cdot 2pq} \tag{1}$$

$$\mathbb{E}_{\Delta X}\left[\sum_{i=1}^{n} h_i^2\right] = \frac{1+3pq}{n \cdot 2pq} \tag{2}$$

$$\mathbb{E}\left[\frac{\Delta X_i^3}{S_n^2}\right] = 0 \tag{3}$$

This section provides the detailed derivations of **Eqs. 1–3**. Throughout, we assume Hardy-Weinberg equilibrium (HWE) and Mendelian segregation.

#### S1.1 Derivation of $\mathrm{Var}_{\Delta X}[\Delta X_i] = 2pq$

As a prerequisite for **Eqs. 1–3**, we first show that the variance of the within-family genotype difference is

$$\mathrm{Var}[\Delta X_i] = 2pq, \quad i = 1, \dots, n. \tag{4}$$

We assume that the focal SNP has minor allele frequency $p$ (with $q = 1 - p$) in a population under Hardy-Weinberg equilibrium (HWE), with Mendelian segregation. Let $X_{j,i} \in \{0, 1, 2\}$ denote the genotype of sibling $j \in \{1, 2\}$ in family $i \in \{1, \dots, n\}$. Each genotype is the sum of the paternal and maternal transmitted alleles:

$$X_{j,i} = F_{j,i} + M_{j,i},$$

where $F_{j,i}$ and $M_{j,i}$ denote the alleles transmitted from the father and mother, respectively. Marginally, each transmitted allele follows a Bernoulli distribution with success probability $p$ under Mendelian segregation. The within-family genotype difference decomposes by parental origin:

$$\begin{aligned}\Delta X_i &= X_{1,i} - X_{2,i} \\ &= (F_{1,i} + M_{1,i}) - (F_{2,i} + M_{2,i}) \\ &= F_{1,i} - F_{2,i} + M_{1,i} - M_{2,i} \\ &:= \Delta F_i + \Delta M_i.\end{aligned} \tag{5}$$

By **Eq. 5** and independent transmissions from the father and the mother ($\Delta F_i \perp\!\!\!\perp \Delta M_i$) under Mendelian segregation, the variance decomposes as:

$$\begin{aligned}\mathrm{Var}\left[\Delta X_i\right] &= \mathrm{Var}\left[\Delta F_i\right] + \mathrm{Var}\left[\Delta M_i\right] \\ &= 2\mathrm{Var}\left[\Delta F_i\right].\end{aligned} \tag{6}$$

We now calculate $\mathrm{Var}[\Delta F_i]$:

$$\mathrm{Var}\left[\Delta F_i\right] = \mathrm{Var}\left[F_{1,i} - F_{2,i}\right] = \mathrm{Var}\left[F_{1,i}\right] + \mathrm{Var}\left[F_{2,i}\right] - 2\mathrm{Cov}\left[F_{1,i}, F_{2,i}\right]. \tag{7}$$

Because each transmitted allele follows a Bernoulli distribution with probability $p$, its variance is then $pq$:

$$\mathrm{Var}[F_{j,i}] = pq, \quad j = 1, 2.$$

We now calculate the covariance term in **Eq. 7**. By the law of total covariance,

$$\begin{aligned}
\mathrm{Cov}\left[F_{1,i}, F_{2,i}\right] &= \mathbb{E}\left[\mathrm{Cov}[F_{1,i}, F_{2,i} \mid G_i]\right] + \mathrm{Cov}\left[\mathbb{E}[F_{1,i} \mid G_i], \mathbb{E}[F_{2,i} \mid G_i]\right] \\
&= 0 + \mathrm{Cov}\left[\frac{G_i}{2}, \frac{G_i}{2}\right] \\
&= \frac{1}{4}\mathrm{Var}\left[G_i\right] \\
&= \frac{1}{2}pq.
\end{aligned}$$

Substituting $\mathrm{Var}\left[F_{j,i}\right] = pq$ and $\mathrm{Cov}\left[F_{1,i}, F_{2,i}\right] = \frac{1}{2}pq$ into **Eq. 7**, the variance of the difference in the transmitted alleles is:

$$\mathrm{Var}\left[\Delta F_i\right] = pq + pq - 2 \cdot \frac{1}{2}pq = pq. \tag{8}$$

Finally, substituting **Eq. 8** back into **Eq. 6** gives the variance of the sibling genotype difference:

$$\mathrm{Var}\left[\Delta X_i\right] = 2pq, \quad i = 1, \ldots, n. \tag{9}$$

#### S1.2 Derivation of $\mathbb{E}\left[\frac{1}{S_n}\right] = \frac{1}{n \cdot 2pq}$

In the main text, we defined $S_n = \sum_{i=1}^{n} \Delta X_i^2$. To derive the expectation of $\frac{1}{S_n}$, we expand $g(S_n) = \frac{1}{S_n}$ by its second-order Taylor expansion:

$$\begin{aligned}
g(S_n) &\approx g\left(\mathbb{E}_{\Delta X}\left[S_n\right]\right) + g'(\mathbb{E}_{\Delta X}[S_n]) \cdot (S_n - \mathbb{E}_{\Delta X}[S_n]) + \frac{1}{2}g''\left(\mathbb{E}_{\Delta X}[S_n]\right)(S_n - \mathbb{E}_{\Delta X}[S_n])^2 \\
&= \frac{1}{\mathbb{E}_{\Delta X}[S_n]} + g'(\mathbb{E}_{\Delta X}[S_n]) \cdot (S_n - \mathbb{E}_{\Delta X}[S_n]) + \left(\frac{1}{\mathbb{E}_{\Delta X}[S_n]^3}\right) \cdot (S_n - \mathbb{E}_{\Delta X}[S_n])^2.
\end{aligned} \tag{10}$$

Taking expectations on both sides of **Eq. 10** yields:

$$\begin{aligned}
\mathbb{E}\left[g(S_n)\right] &= \frac{1}{\mathbb{E}_{\Delta X}[S_n]} + g'(\mathbb{E}_{\Delta X}[S_n]) \cdot (\mathbb{E}_{\Delta X}[S_n] - \mathbb{E}_{\Delta X}[S_n]) + \left(\frac{1}{\mathbb{E}_{\Delta X}[S_n]^3}\right) \cdot \mathrm{Var}[S_n] \\
&= \frac{1}{\mathbb{E}_{\Delta X}[S_n]} + 0 + \frac{\mathrm{Var}_{\Delta X}[S_n]}{\mathbb{E}_{\Delta X}[S_n]^3}.
\end{aligned} \tag{11}$$

We now calculate $\mathbb{E}_{\Delta X}[S_n]$. Because

$$\mathbb{E}[\Delta X_i] = \mathbb{E}[X_{1,i} - X_{2,i}] = \mathbb{E}[X_{1,i}] - \mathbb{E}[X_{2,i}] = 0 \text{ and } \mathrm{Var}[\Delta X_i] = 2pq, i = 1, \ldots, n,$$

the expectation of the squared genotype difference is

$$\mathbb{E}[\Delta X_i^2] = \mathrm{Var}[\Delta X_i] + \mathbb{E}[\Delta X_i]^2 = 2pq.$$

Summing over sibships,

$$\mathbb{E}_{\Delta X}[S_n] = \mathbb{E}_{\Delta X}\left[\sum_{i=1}^{n} \Delta X_i^2\right] = \sum_{i=1}^{n} \mathbb{E}[\Delta X_i^2] = n \cdot 2pq.$$

Because sibling pairs are independent across families, the variance of $S_n$ is:

$$\mathrm{Var}_{\Delta X}[S_n] = \mathrm{Var}\left[\sum_{i=1}^{n} \Delta X_i^2\right] = \sum_{i=1}^{n} \mathrm{Var}[\Delta X_i^2] = n \cdot \mathrm{Var}\left[\Delta X_i^2\right].$$

Because $\mathrm{Var}[\Delta X_i^2]$ depends only on the allele frequency $p$ and not on $n$, it is a constant with respect to the sample size $n$. Therefore,

$$\mathrm{Var}_{\Delta X}[S_n] = O(n).$$

Substituting $\mathbb{E}_{\Delta X}[S_n] = n \cdot 2pq$ and $\mathrm{Var}_{\Delta X}[S_n] = O(n)$ into **Eq. 11** gives the expectation of $\frac{1}{S_n}$:

$$\begin{aligned}\mathbb{E}\left[\frac{1}{S_n}\right] &= \frac{1}{n \cdot 2pq} + \frac{O(n)}{(n \cdot 2pq)^3} \\ &= \frac{1}{n \cdot 2pq} + O(n^{-2}).\end{aligned}$$

#### S1.3 Derivation of $\mathbb{E}_{\Delta X}\left[\sum_{i=1}^{n} h_i^2\right] = \frac{1+3pq}{n \cdot 2pq}$

In the main text, we defined the leverage as:

$$h_i = \frac{\Delta X_i^2}{S_n}, \text{ where } S_n = \sum_{i=1}^{n} \Delta X_i^2.$$

In this subsection, we provide the detailed derivation for the expectation of the sum of squared leverages,

$$\mathbb{E}_{\Delta X}\left[\sum_{i=1}^{n} h_i^2\right] = \frac{1+3pq}{n \cdot 2pq}.$$

Because sibling pairs are independent and identically distributed across families, the expectation of the sum reduces to $n$ times the expectation of a single squared leverage,

$$\mathbb{E}_{\Delta X}\left[\sum_{i=1}^{n} h_i^2\right] = \sum_{i=1}^{n} \mathbb{E}_{\Delta X}[h_i^2] = n \cdot \mathbb{E}_{\Delta X}[h_i^2].$$

We therefore focus on evaluating $\mathbb{E}_{\Delta X}[h_i^2]$.

By definition:

$$h_i^2 = \frac{\Delta X_i^4}{(\sum_{j=1}^{n} \Delta X_j^2)^2} = \frac{\Delta X_i^4}{n^2 \left(\overline{\Delta X^2}\right)^2}, \quad \overline{\Delta X^2} = \frac{\sum_{j=1}^{n} \Delta X_j^2}{n}. \tag{12}$$

We begin by approximating the denominator using the Central Limit Theorem. Since $\Delta X_i^2$ are independent and identically distributed with finite variance, the sample mean satisfies

$$\overline{\Delta X^2} = 2pq + O_p(n^{-\frac{1}{2}}).$$

Therefore,

$$\frac{1}{\left(\overline{\Delta X^2}\right)^2} = \frac{1}{(2pq)^2} \cdot \left(1 + O_p(n^{-\frac{1}{2}})\right).$$

Substituting this approximation back into the squared leverage (**Eq. 12**) gives:

$$h_i^2 = \frac{\Delta X_i^4}{n^2 \cdot (2pq)^2} + O_p(n^{-\frac{5}{2}}). \tag{13}$$

Taking expectations over both sides of **Eq. 13** yields:

$$\mathbb{E}\left[h_i^2\right] = \frac{\mathbb{E}\left[\Delta X_i^4\right]}{n^2(2pq)^2} + O(n^{-3}). \tag{14}$$

To evaluate **Eq. 14**, we next calculate $\mathbb{E}[\Delta X_i^4]$. Using the parental decomposition of $\Delta X_i$ (**Eq. 5**), the expectation of $\Delta X_i^4$ is:

$$\begin{aligned}
\mathbb{E}\left[\Delta X_i^4\right] &= \mathbb{E}\left[(\Delta F_i + \Delta M_i)^4\right] \\
&= \mathbb{E}\left[\Delta F_i^4 + 4\Delta F_i^3\Delta M_i + 6\Delta F_i^2\Delta M_i^2 + 4\Delta F_i\Delta M_i^3 + \Delta M_i^4\right] \\
&= \mathbb{E}\left[\Delta F_i^4\right] + 4\mathbb{E}\left[\Delta F_i^3\Delta M_i\right] + 6\mathbb{E}\left[\Delta F_i^2\Delta M_i^2\right] + 4\mathbb{E}\left[\Delta F_i\Delta M_i^3\right] + \mathbb{E}\left[\Delta M_i^4\right].
\end{aligned} \tag{15}$$

The odd-order cross terms in **Eq. 15** vanish because $\Delta F_i$ and $\Delta M_i$ are independent with mean zero; then $\mathbb{E}[\Delta F_i^3\Delta M_i] = \mathbb{E}[\Delta F_i^3]\mathbb{E}[\Delta M_i] = 0$. In addition, $\Delta F_i$ and $\Delta M_i$ are independent and identically distributed under random mating and Mendelian segregation. **Eq. 15** simplifies to:

$$\mathbb{E}\left[\Delta X_i^4\right] = 2\mathbb{E}\left[\Delta F_i^4\right] + 6\mathbb{E}\left[\Delta F_i^2\right]^2. \tag{16}$$

We obtain $\mathbb{E}[\Delta F_i^2]$ and $\mathbb{E}[\Delta F_i^4]$ from the distribution of $\Delta F_i$. The random variable $\Delta F_i$ takes values $\{-1, 0, 1\}$ with probabilities:

$$\begin{aligned}
\mathbb{P}\left[\Delta F_i = 1\right] &= \sum_{G_i} \mathbb{P}\left[\Delta F_i = 1 \mid G_i\right] = 0 \cdot p^2 + \frac{1}{4} \cdot 2pq + 0 \cdot q^2 = \frac{1}{2}pq, \\
\mathbb{P}\left[\Delta F_i = -1\right] &= \sum_{G_i} \mathbb{P}\left[\Delta F_i = -1 \mid G_i\right] = \frac{1}{2}pq, \\
\mathbb{P}\left[\Delta F_i = 0\right] &= \sum_{G_i} \mathbb{P}\left[\Delta F_i = 0 \mid G_i\right] = 1 \cdot p^2 + \frac{1}{2} \cdot 2pq + 1 \cdot q^2 = p^2 + q^2 + pq.
\end{aligned}$$

Hence, the expectations of $\Delta F_i^2$ and $\Delta F_i^4$ are:

$$\mathbb{E}\left[\Delta F_i^2\right] = \sum_{\Delta f_i} \Delta f_i^2 \cdot \mathbb{P}\left[\Delta F_i = \Delta f_i\right] = 1^2 \cdot \frac{1}{2}pq + (-1)^2 \cdot \frac{1}{2}pq + 0^2 \cdot (p^2 + q^2 + pq) = pq. \tag{17}$$

$$\mathbb{E}\left[\Delta F_i^4\right] = \sum_{\Delta f_i} \Delta f_i^4 \cdot \mathbb{P}\left[\Delta F_i = \Delta f_i\right] = pq. \tag{18}$$

Substituting **Eq. 17** and **Eq. 18** into **Eq. 16** gives the expectation of $\Delta X_i^4$:

$$\mathbb{E}\left[\Delta X_i^4\right] = 2pq + 6(pq)^2.$$

Substituting this result back into **Eq. 14**, the expectation of the squared leverage is:

$$\begin{aligned}\mathbb{E}\left[h_i^2\right] &= \frac{2pq + 6(pq)^2}{n^2(2pq)^2} + O(n^{-3}) \\ &= \frac{1 + 3pq}{n^2 \cdot 2pq} + O(n^{-3}). \end{aligned} \tag{19}$$

Finally, the expectation of the sum over the squared leverages is:

$$\mathbb{E}_{\Delta X}\left[\sum_{i=1}^{n} h_i^2\right] = \frac{1 + 3pq}{n \cdot 2pq} + O(n^{-2}).$$

#### S1.4 Derivation of $\mathbb{E}\left[\frac{\Delta X_i}{S_n}\right] = \mathbb{E}\left[\frac{\Delta X_i^3}{S_n}\right] = \mathbb{E}\left[\frac{\Delta X_i^3}{S_n^2}\right] = 0$

The key idea underlying these results is that the distribution of $\Delta X_i$ (and $\Delta X_i^3$) is symmetric around 0, while any power of $S_n$ depends on $\Delta X_i$ only through $\Delta X_i^2$. As a result, $\frac{\Delta X_i}{S_n}$, $\frac{\Delta X_i^3}{S_n}$, and $\frac{\Delta X_i^3}{S_n^2}$ are odd functions of $\Delta X_i$ and their expectations are zero. To see this, we write $S_n$ as a sum

$$\begin{aligned} S_n &= \Delta X_i^2 + \sum_{j \neq i} \Delta X_j^2 \\ &= \Delta X_i^2 + T_i, \end{aligned}$$

where $T_i := \sum_{j \neq i} \Delta X_j^2$. Because sibling pairs are independent across families, $\Delta X_i$ is independent of $T_i$. Therefore, conditional on $T_i$, the only random term in the ratio is $\Delta X_i$. For fixed $T_i$, $\frac{\Delta X_i}{\Delta X_i^2 + T_i}$, $\frac{\Delta X_i^3}{(\Delta X_i^2 + T_i)^2}$, and $\frac{\Delta X_i^3}{\Delta X_i^2 + T_i}$ are each odd functions of $\Delta X_i$. Moreover, the distributions of $\Delta X_i$ (and $\Delta X_i^3$) are symmetric around 0. The conditional expectations are well-defined because, under Hardy-Weinberg equilibrium with $0 < p < 1$, the denominator $S_n > 0$ with probability $1 - (1 - pq)^{2n} > 0$. Therefore, all three conditional expectations are zero:

$$\begin{aligned} \mathbb{E}\left[\frac{\Delta X_i}{\Delta X_i^2 + T_i} \mid T_i\right] &= 0, \\ \mathbb{E}\left[\frac{\Delta X_i^3}{\Delta X_i^2 + T_i} \mid T_i\right] &= 0, \\ \mathbb{E}\left[\frac{\Delta X_i^3}{(\Delta X_i^2 + T_i)^2} \mid T_i\right] &= 0. \end{aligned}$$

By the law of total expectation, the expectation of $\frac{\Delta X_i}{S_n}$ and $\frac{\Delta X_i^3}{S_n}$ is zero:

$$\mathbb{E}\left[\frac{\Delta X_i}{S_n}\right] = \mathbb{E}_{T_i}\left[\mathbb{E}_{\Delta X_i}\left[\frac{\Delta X_i}{\Delta X_i^2 + T_i} \mid T_i\right]\right] = 0,$$
$$\mathbb{E}\left[\frac{\Delta X_i^3}{S_n}\right] = \mathbb{E}_{T_i}\left[\mathbb{E}_{\Delta X_i}\left[\frac{\Delta X_i^3}{\Delta X_i^2 + T_i} \mid T_i\right]\right] = 0,$$
$$\mathbb{E}\left[\frac{\Delta X_i^3}{S_n^2}\right] = \mathbb{E}_{T_i}\left[\mathbb{E}_{\Delta X_i}\left[\frac{\Delta X_i^3}{(\Delta X_i^2 + T_i)^2} \mid T_i\right]\right] = 0.$$

### S2 Lemmas used to derive the bias of the block jackknife estimator.

In the **Methods** , we used the following results to derive the bias of the block jackknife variance estimator.

$$\mathrm{Var}[Q_n] = O(n^{-2}),$$
$$\mathrm{Var}_I\left[\frac{1}{2}(\bar{W}_{(m)} - \mu)^T H(\bar{W}_{(m)} - \mu)\right] = O(r^{-2}),$$
$$\mathrm{Cov}_I\left[\bar{\psi}_{(m)}, \frac{1}{2}(\bar{W}_{(m)} - \mu)^T H(\bar{W}_{(m)} - \mu) \mid W\right] = O(r^{-3/2}).$$

In this section, we provide the detailed derivations of these results.

#### S2.1 $\mathrm{Var}[Q_n]$ is of order $O(n^{-2})$.

$Q_n$ is the second order term in the Taylor expansion for $\hat{\beta}_0 - \beta_0$:

$$Q_n = \frac{1}{2}\left(\bar{W}_n - \mu\right)^T H\left(\bar{W}_n - \mu\right), \quad \bar{W}_n = \frac{\sum_{i=1}^n W_i}{n}, \; H = \nabla^2 g(\mu). \tag{20}$$

By the Central Limit Theorem,

$$\sqrt{n}(\bar{W}_n - \mu) \xrightarrow{d} \mathcal{N}(0, \Sigma).$$

We write $Z = \sqrt{n}(\bar{W}_n - \mu) \xrightarrow{d} \mathcal{N}(0, \Sigma)$. The second-order term $Q_n$ can then be rewritten as

$$\begin{aligned} Q_n &= \frac{1}{2}(\bar{W}_n - \mu)^T H(\bar{W}_n - \mu) \\ &= \frac{1}{2}\left(\frac{1}{\sqrt{n}}Z\right)^T H\left(\frac{1}{\sqrt{n}}Z\right) \\ &= \frac{1}{2n}Z^T HZ. \end{aligned}$$

Because $H = \nabla^2 g(\mu)$ is symmetric, $Q_n = \frac{1}{2n} Z^T H\, Z$ is a Gaussian quadratic form whose variance is

$$\mathrm{Var}[Q_n] = \frac{1}{2n^2}\mathrm{tr}\left[H\Sigma H\Sigma\right]. \tag{21}$$

We next show that $\mathrm{tr}[H\Sigma H\Sigma]$ is a finite constant, so that $\mathrm{Var}[Q_n] = O(n^{-2})$.

By the definition of $H = \nabla^2 g(\mu)$ and $g(u,v) = \frac{u}{v}$,

$$H = \nabla^2 g(u,v) = \begin{bmatrix} \frac{\partial^2 g}{\partial u^2}(u,v) & \frac{\partial^2 g}{\partial u\,\partial v}(u,v) \\ \frac{\partial^2 g}{\partial v\,\partial u}(u,v) & \frac{\partial^2 g}{\partial v^2}(u,v) \end{bmatrix}_{(u,v)=(\beta_0\mu_2,\mu_2)}$$
$$= \begin{bmatrix} 0 & -v^{-2} \\ -v^{-2} & 2u\,v^{-3} \end{bmatrix}_{(u,v)=(\beta_0\mu_2,\mu_2)}$$
$$= \begin{bmatrix} 0 & -\mu_2^{-2} \\ -\mu_2^{-2} & 2\beta_0\,\mu_2^{-2} \end{bmatrix}. \tag{22}$$

$\Sigma$ is the covariance matrix of the observed vector $W_i = \begin{bmatrix} \Delta X_i \cdot \Delta Y_i \\ \Delta X_i^2 \end{bmatrix}$:

$$\Sigma = \mathrm{Cov}\begin{bmatrix} \Delta X_i \Delta Y_i \\ \Delta X_i^2 \end{bmatrix} = \begin{bmatrix} \mathrm{Var}[\Delta X_i \cdot \Delta Y_i] & \mathrm{Cov}[\Delta X_i \cdot \Delta Y_i, \Delta X_i^2] \\ \mathrm{Cov}[\Delta X_i \cdot \Delta Y_i, \Delta X_i^2] & \mathrm{Var}[\Delta X_i^2] \end{bmatrix}.$$

Denoting $\mathbb{E}[\Delta X_i^k] = \mu_k$, the covariance matrix $\Sigma$ is:

$$\Sigma = \begin{bmatrix} \beta_0^2(\mu_4 - \mu_2^2) + \tau^2\mu_4 + \mathbb{E}_{\Delta X}[\Delta X_i^2 \sigma^2(\Delta X_i^2)] & \beta_0(\mu_4 - \mu_2^2) \\ \beta_0(\mu_4 - \mu_2^2) & \mu_4 - \mu_2^2 \end{bmatrix}. \tag{23}$$

Using **Eq. 22** and **Eq. 23**, the product of the two matrices $H$ and $\Sigma$ is:

$$H\Sigma = \begin{bmatrix} 0 & -\mu_2^{-2} \\ -\mu_2^{-2} & 2\beta_0\mu_2^{-2} \end{bmatrix} \cdot \begin{bmatrix} \beta_0^2(\mu_4 - \mu_2^2) + \tau^2\mu_4 + \mathbb{E}_{\Delta X}[\Delta X_i^2 \sigma^2(\Delta X_i^2)] & \beta_0(\mu_4 - \mu_2^2) \\ \beta_0(\mu_4 - \mu_2^2) & \mu_4 - \mu_2^2 \end{bmatrix}$$
$$= \begin{bmatrix} -\beta_0\mu_2^{-2}(\mu_4 - \mu_2^2) & -\mu_2^{-2}(\mu_4 - \mu_2^2) \\ -\beta_0^2\mu_2^{-2}(\mu_4 - \mu_2^2) - \tau^2\mu_2^{-2}\mu_4 - \mu_2^{-2}\mathbb{E}_{\Delta X}[\Delta X_i^2\sigma^2(\Delta X_i^2)] + 2\beta_0^2\mu_2^{-2}(\mu_4 - \mu_2^2) & \beta_0\mu_2^{-2}(\mu_4 - \mu_2^2) \end{bmatrix}.$$

For any $2 \times 2$ matrix $A = \begin{pmatrix} a & b \\ c & d \end{pmatrix}$, the trace of its square $A^2$ is $a^2 + 2bc + d^2$. Therefore, the trace of $H\Sigma H\Sigma$ is:

$$\mathrm{tr}[H\Sigma H\Sigma] = \frac{2}{\mu_2^4}\,(\mu_4 - \mu_2^2)\left(\tau^2\mu_4 + \mathbb{E}_{\Delta X}\left[\sigma^2(\Delta X_i^2)\,\Delta X_i^2\right]\right) =: C_2 < \infty. \tag{24}$$

As a result, $\mathrm{Var}[Q_n] = \frac{1}{2n^2} C_2 = O(n^{-2})$, confirming that the second-order contribution to the variance is of order $O(n^{-2})$.

#### S2.2 The variance of the second-order terms in the Taylor expansion of the block jackknife estimator.

In this section, we show that the second-order variance term $\mathrm{Var}_I\left[\frac{1}{2}(\bar{W}_{(m)} - \mu)^T H\,(\bar{W}_{(m)} - \mu) \mid W\right]$ and the cross term $\mathrm{Cov}_I\left[\bar{\psi}_{(m)}, \frac{1}{2}(\bar{W}_{(m)} - \mu)^T H\,(\bar{W}_{(m)} - \mu) \mid W\right]$ are of order $O(r^{-2})$ and $O(r^{-3/2})$, respectively—both negligible relative to the leading $O(r^{-1})$ contribution from $\mathrm{Var}_I[\bar{\psi}_{(m)} \mid W]$.

As with the full-sample case, define $Z_{(m)} = \sqrt{r}\,(\bar{W}_{(m)} - \mu)$. Because $W_i$ have bounded fourth moments (under our model assumptions), $Z_{(m)}$ has a covariance matrix with bounded entries, and the quadratic form $Z_{(m)}^T H\, Z_{(m)}$ has bounded variance. Since the Hessian matrix $H$ and the population covariance $\Sigma$ are fixed, it follows that

$$\mathrm{Var}_I\left[\frac{Z_{(m)}^T H\, Z_{(m)}}{2r} \,\middle|\, W\right] = \frac{1}{4r^2}\,\mathrm{Var}\Big[Z_{(m)}^T H\, Z_{(m)}\Big] = O(r^{-2}). \tag{25}$$

For the covariance term, $\mathrm{Cov}_I\Big[\bar{\psi}_{(m)},\ \frac{1}{2}(\bar{W}_{(m)} - \mu)^T H\,(\bar{W}_{(m)} - \mu) \mid W\Big]$, we apply the Cauchy–Schwarz inequality:

$$\begin{aligned}
\Big|\mathrm{Cov}_I\Big[\bar{\psi}_{(m)},\ \tfrac{1}{2}(\bar{W}_{(m)} - \mu)^T H\,(\bar{W}_{(m)} - \mu) \mid W\Big]\Big| &\le \sqrt{\mathrm{Var}_I\Big[\bar{\psi}_{(m)} \mid W\Big] \cdot \mathrm{Var}_I\Big[\tfrac{1}{2}(\bar{W}_{(m)} - \mu)^T H\,(\bar{W}_{(m)} - \mu) \,\Big|\, W\Big]} \\
&= \sqrt{O(r^{-1}) \cdot O(r^{-2})} \\
&= O(r^{-3/2}).
\end{aligned}$$

### S3 A simulation study validates the analytical derivation results

In the **Results** section of the main text, we derived the biases of three estimators for the variance of allelic effect estimates (**Eq. 7**, **Eq. 8**, and **Eq. 9**). Here, we validated these derivations through simulation. Our simulations examined how the bias of each variance estimator changed with respect to the key parameters—the baseline allelic effect ($\beta_0$), the variance of allelic effects ($\tau^2$), the heteroskedasticity factor ($\tau^2 + \sigma_1$), sample size ($n$), and minor allele frequency (maf).

We simulated a population of 5,000 sibling pairs and one focal SNP. The parental genotypes at the focal SNP were drawn from Hardy-Weinberg equilibrium at a maf of 10%. Allele transmission followed Mendelian segregation. Individual phenotypes were sampled according to the individual-level model (**Eq. 1**), wherein phenotype values were the sum of a baseline phenotype, a focal SNP effect, and a non-focal random effect. Siblings had correlated non-focal effects reflecting shared genetic background and environment. Therefore, the non-focal variation of each sibling pair was drawn from a family-specific bivariate normal distribution with non-zero covariance. Following the heteroskedasticity assumption in the main text (**Eq. 4**), the variance of the non-focal term depended on the focal genotype contrast of each sibling pair. Specifically, the non-focal variance of the $i$-th family changed linearly with the genotype contrast:

$$\mathrm{Var}\left[\Delta\epsilon_i\right] = \sigma_0 + \sigma_1 \cdot \Delta X_i^2.$$

Building upon this simulation design, we conducted three distinct sets of simulations (corresponding to the three rows in **Fig. S1**) under varying conditions. Across all scenarios, to account for Monte Carlo noise, we generated 5,000 independent simulation replicates using different random seeds for each parameter set. In all corresponding figures, the mean variance estimates across the 5,000 independent simulation replicates minus the derived theoretical variance (**Eq. 5**) are plotted as dots. These align with our theoretical predictions, plotted as solid lines.

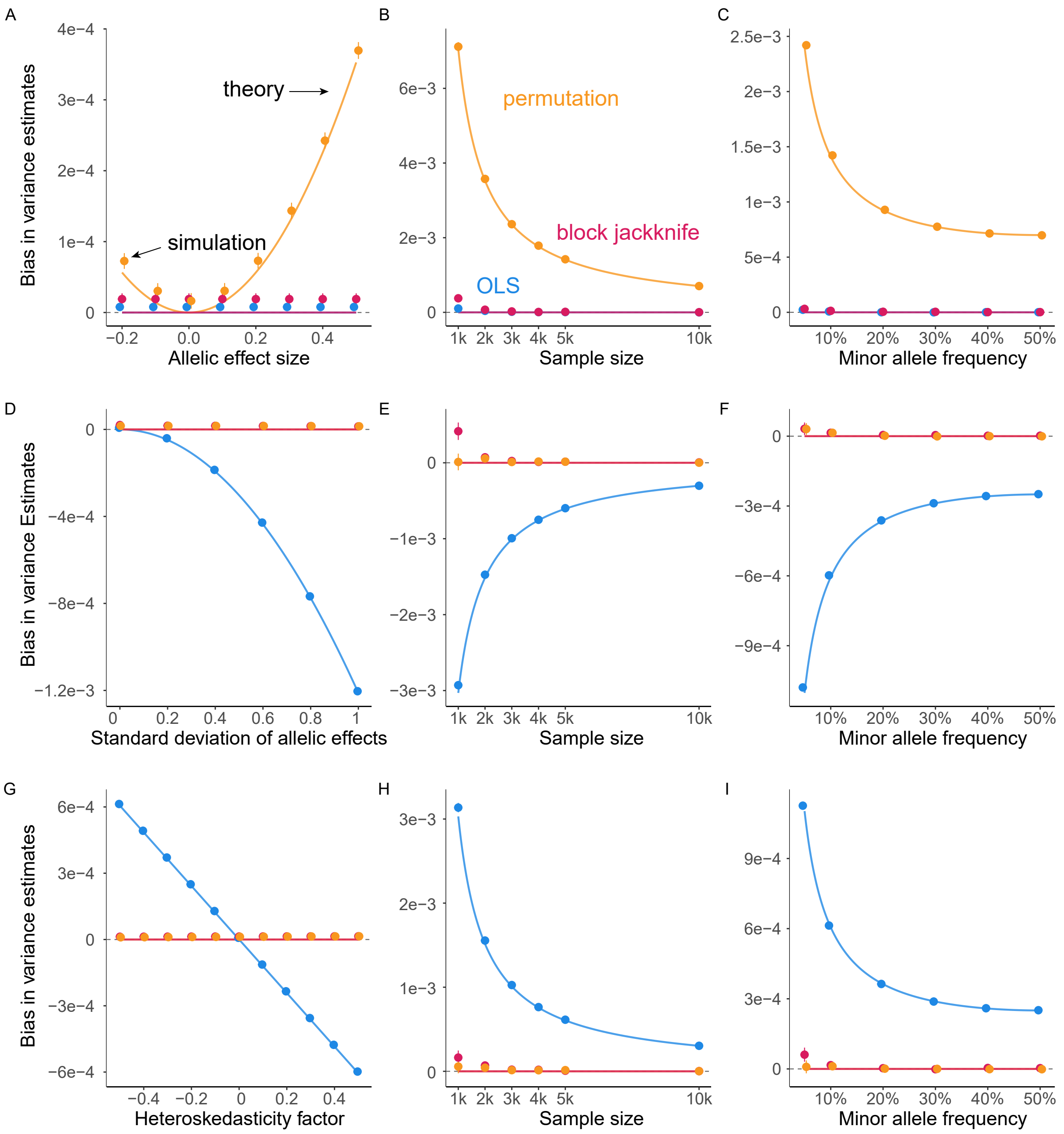

**Figure S1. Simulations validate theoretical derivations.** Markers show the estimated bias of the variance estimators: the mean variance estimates across 5000 simulation replicates minus the true variance (**Eq. 5** in the main text). Error bars show the mean $\pm 1.96 \cdot$SE of the estimated bias. Lines show the derivation results from **Eqs. 7, 8, 9** in the main text. Unless otherwise specified, each simulation included 5000 pairs of siblings, with parental genotypes at the focal SNP drawn from the Hardy-Weinberg equilibrium at a maf of 10%, a baseline allelic effect of zero, and a heteroskedasticity factor of zero. **(A)** varies the baseline allelic effects. **(B)** and **(C)** vary sample size and maf, respectively, with the baseline allelic effect set to 1. **(D)** varies the standard deviation of allelic effects. **(E)** and **(F)** vary sample size and maf, respectively, with the variance of the allelic effects set to 0.5. **(G)** varies the heteroskedasticity factor. **(H)** and **(I)** vary sample size and maf, respectively, with the heteroskedasticity factor set to -0.5.

**Non-zero baseline allelic effects.** First, we examined bias by varying the baseline allelic effect $\beta_0$ from $-0.2$ to 0.5 in increments of 0.1, keeping the variance of allelic effects and heteroskedasticity factor at 0 (**Fig. S1A**). Next, we fixed the baseline effect at 1 and evaluated the estimators across varying sample sizes (1k, 2k, 3k, 4k, 5k, and 10k sibships) (**Fig. S1B**) and mafs (5%, 10%, 20%, 30%, 40%, 50%) (**Fig. S1C**).

**Non-zero variance of allelic effects.** In a similar design, we varied the standard deviation of the allelic effects from 0 to 1 in increments of 0.2, keeping the baseline allelic effect at zero and the non-focal variation homoskedastic (**Fig. S1D**). We then fixed the variance of the allelic effects at 0.5, evaluating across the same varying sample sizes (**Fig. S1E**) and mafs (**Fig. S1F**).

**Non-zero heteroskedasticity factor.** Lastly, we varied the heteroskedasticity factor (here equal to $\sigma_1$, since $\tau^2 = 0$) from $-0.5$ to 0.5 in increments of 0.1, keeping the baseline allelic effect and its variance at 0 (**Fig. S1G**). Next, we fixed the heteroskedasticity factor at $-0.5$ and evaluated the estimators across the same varying sample sizes (**Fig. S1H**) and mafs (**Fig. S1I**).

Overall, our simulations show high concordance with our theoretical derivations in the **Results** section and thereby serve to validate them.

### S4 Rare alleles

Accurate uncertainty quantification for rare alleles is of particular biological interest because they tend to have larger effects (due to stabilizing or purifying selection)[1–3]. In this section, we examine the sampling distribution of the variance estimators for rare alleles. We then investigate the reason for the apparent discrepancy between our prediction of a positive bias for the permutation estimator and the fact that in UKB data we estimate a negative bias for non-significant rare alleles (**Figs. 3, S9**).

#### S4.1 Skewed distributions of variance estimates for rare alleles

**Model setup.** Following the simulation design described in **Section S3**, we simulated 5,000 sibling pairs. Parental genotypes at the focal SNP were sampled from the Hardy-Weinberg equilibrium at maf 1%, and offspring genotypes were generated by Mendelian transmission. The allelic effect was fixed at 0, and the non-focal variation was homoskedastic across families. Under these conditions, the variance estimators were unbiased. Here, we examined the distribution of each variance estimator by computing the difference between the estimated variance and the true variance of allelic effect estimates (**Eq. 5**).

**Simulations and results.** All four variance estimators—OLS, permutation, block jackknife and the cluster-based method (introduced in **Section S5**)—exhibited negligible bias on average (**Fig. S2**), consistent with our conclusions in the main text. However, their sampling distributions are substantially right-skewed. This skewness arises from the sharp decrease in effective sample size for rare alleles. When the allele is rare, most families are homozygous at the focal

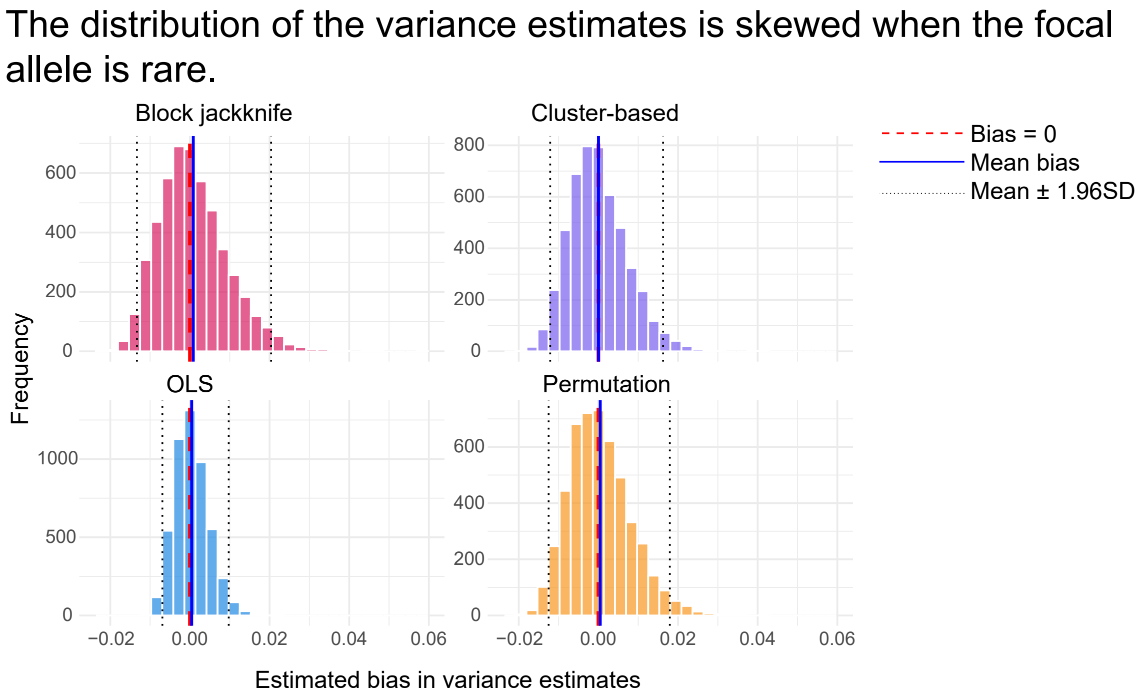

**Figure S2. The distribution of the variance estimators is skewed when the focal allele is rare.** The x-axis shows the estimated bias in the four variance estimators, calculated as the variance estimates minus the derived variance (**Eq. 5**) for each of the 5,000 simulation replicates. Each simulation included 5000 sibling pairs with the parental genotypes at the focal SNP sampled from the Hardy-Weinberg equilibrium at maf 1%. The true allelic effect was fixed at 0, and the non-focal variation was homoskedastic across families. While the variance estimators remain approximately unbiased, their sampling distributions are right-skewed under low minor allele frequency.

locus and contribute little information about the allelic effect. Because block jackknife, cluster-based, and permutation variance estimators are constructed from the sums of squared deviations of approximately normally distributed quantities, their sampling distributions approximate a $\chi^2$-distribution. The OLS estimator is a function of the residual sum of squares, and its sampling distribution approximates a $\chi^2$-distribution. Because maf is low, few sibling pairs provide informative comparisons, and therefore, the effective sample size is low. The $\chi^2$-distribution has low degrees of freedom and is therefore right-skewed[4] (**Fig. S2**).

#### S4.2 Finite-sample bias of the ratio between permutation and block jackknife variance estimates under rare alleles

In **Fig. 3C** and **Fig. S9**, we observed that for non-significant SNPs ($p > 0.05$) with low minor allele frequency, the permutation variance estimates were smaller than the block jackknife benchmark, a direction opposite to the theoretical prediction that the permutation bias is non-negative (**Eq. 8**). We hypothesized that the discrepancy reflects a finite-sample property of the ratio-based statistic itself, rather than a violation of our primary results. Specifically, the $\log_2$ ratio is a nonlinear function of two correlated variance estimators, and its expectation depends on the higher-order moments of their joint distribution. These higher-order contributions are negligible when the effective sample size is large but become pronounced when the effective sample size is small, as occurs for rare alleles. To test this hypothesis, we conducted simulations across a range of low minor allele frequencies and performed a downsampling analysis of the empirical data. Results are shown for diastolic blood pressure, which was the representative trait in **Fig. 3**; similar patterns were observed across all 17 traits.

**Support from simulations.** Following the simulation design described in **Section S3**, we simulated 5,000 sibling pairs. Parental genotypes were sampled from the Hardy-Weinberg distribution at maf $\in \{0.1\%, 0.2\%, 0.3\%, 0.4\%, 0.5\%\}$, with offspring genotypes generated by Mendelian transmission. The allelic effect was fixed at zero, and the non-focal variation was homoskedastic across families, so that both permutation and block jackknife variance estimators are unbiased. Under these conditions, the higher-order moments of $\log_2$ ratio are expected to be more pronounced when the maf decreases, and converge to zero when the maf increases.

The simulation results showed that the sample mean of $\log_2(\widehat{\mathrm{Var}}_{perm}[\hat{\beta}_0]/\widehat{\mathrm{Var}}_{bjk}[\hat{\beta}_0])$ fell below zero, and the magnitude of the deviation increased as minor allele frequency decreased (**Fig. S3A**). This pattern is consistent with the $\log_2$ ratio being a biased estimator when the effective sample size is small: at low maf, few sibling pairs are informative about the allelic effect, and the effective sample size is low. With the low effective sample size, the higher-order moments contribute substantially to the expectation of the $\log_2$ ratio statistic.

**Support from empirical downsampling in UKB.** To verify that the same effect operated in the empirical data, we randomly selected subsets of $x$ SNPs, with $x \in \{500, 1000, 5000, 20000\}$ from the $504,858$ SNPs available and showed the results for diastolic blood pressure. Within each subset, we restricted to non-significant ($p \geq 0.05$), rare alleles (maf$< 1\%$) and examined the distribution of $\log_2(\widehat{\mathrm{Var}}_{perm}[\hat{\beta}_0]/\widehat{\mathrm{Var}}_{bjk}[\hat{\beta}_0])$. As predicted, when fewer SNPs were available, the sample mean of $\log_2$ ratio deviated further from zero (**Fig. S3B**). The observation is consistent with the finite-sample bias of the ratio statistic becoming more pronounced when fewer SNPs are available.

Together, the simulation and downsampling results indicate that the downward deviation of the $\log_2(\widehat{\mathrm{Var}}_{perm}[\hat{\beta}_0]/\widehat{\mathrm{Var}}_{bjk}[\hat{\beta}_0])$ estimator of the bias for rare, non-significant SNPs in **Fig. 3** and **Fig. S9** reflects a substantial finite-sample bias of this $\log_2$ ratio itself, rather than a contradiction of our derivation for the bias of the permutation-based estimator. The estimator of the bias works well when the effective sample size is large. However, for rare alleles, where only a few sibling pairs are informative about the allelic effect, the effective sample size is small.

### S5 Performance of allelic effect variance estimators under more complex models

Throughout the main text, we assumed our sample was composed of independent sibships with no between-family covariances or population structure. Here, in light of the increasing availability and interest in analyzing larger and more diverse forms of family data, we revisit these

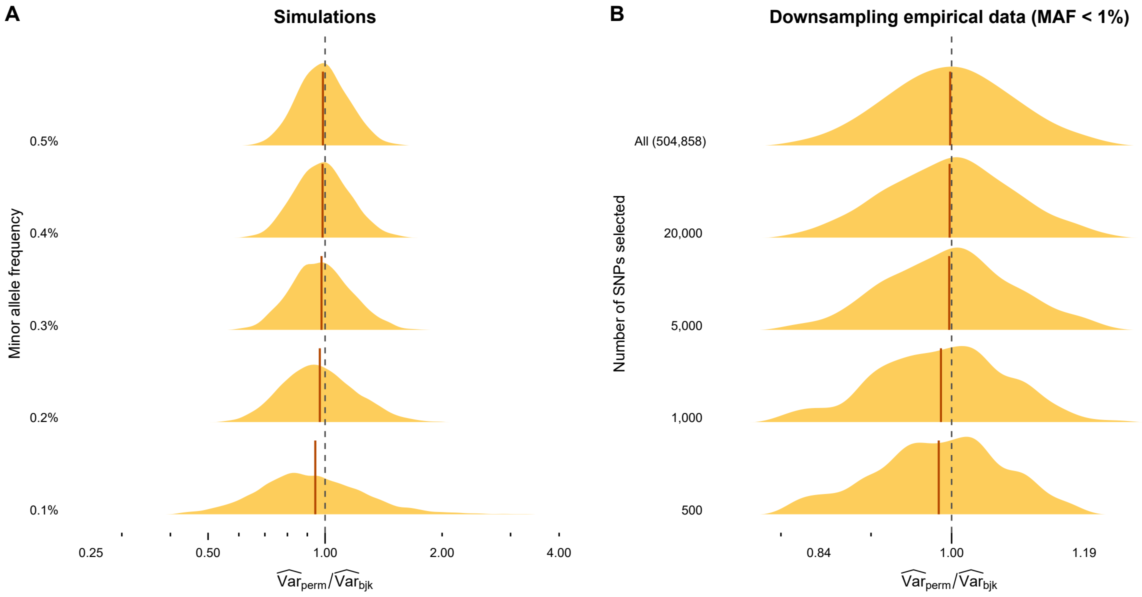

**Figure S3. The sample mean of** $\log_2(\widehat{\mathrm{Var}}_{perm}[\hat{\beta}_0]/\widehat{\mathrm{Var}}_{bjk}[\hat{\beta}_0])$ **deviates from zero when the focal allele is rare.** The estimated $\log_2(\widehat{\mathrm{Var}}_{perm}[\hat{\beta}_0]/\widehat{\mathrm{Var}}_{bjk}[\hat{\beta}_0])$ is used to estimate the bias in the permutation estimator in **Fig. 3**. Solid red lines show the sample mean across replicates, and dashed vertical lines mark zero. **(A)** Distribution of $\log_2(\widehat{\mathrm{Var}}_{perm}[\hat{\beta}_0]/\widehat{\mathrm{Var}}_{bjk}[\hat{\beta}_0])$ across 5,000 simulations across scenarios with different minor allele frequencies (on the y-axis). Each simulation included 5,000 sibling pairs with parental genotypes drawn from the Hardy-Weinberg equilibrium at the indicated maf (on the y-axis). The allelic effect was fixed at zero, and the non-focal variation was homoskedastic across families. **(B)** Distribution of $\log_2(\widehat{\mathrm{Var}}_{perm}[\hat{\beta}_0]/\widehat{\mathrm{Var}}_{bjk}[\hat{\beta}_0])$ from random subsets of the empirical data for diastolic blood pressure. From each random subset, we selected the non-significant ($p > 0.05$), rare (maf < 1%) alleles.

assumptions and show that our main conclusions from the main text extend to more complex scenarios.

Additionally, recent work[5] applied the cluster-based method (also known as the robust sandwich estimator[6]) to allelic effect variance estimation in sib-GWAS by weighting each family's contribution to the variance estimate by its own squared residual rather than assuming a common residual variance across families. For the case where family $i$ contains $n_i$ siblings, let $\tilde{X}_{j,i} = X_{j,i} - \bar{X}_i$ ($\bar{X}_i = \frac{\sum_{j=1}^{n_i} X_{j,i}}{n_i}$) denote the centered genotype of sibling $j$ in family $i$. The cluster-based variance estimator is:

$$\widehat{\text{Var}}_{cluster}[\hat{\beta}] = \frac{n}{n-1} \frac{\sum_{i=1}^{n} \left( \sum_{j=1}^{n_i} \tilde{X}_{j,i}(Y_{j,i} - \hat{Y}_{j,i}) \right)^2}{(\sum_{i=1}^{n} \sum_{j=1}^{n_i} \tilde{X}_{j,i}^2)^2}.$$

Below, we show that the cluster-based method and the block jackknife method yield comparable performance—they are both generally the least biased in the face of the complications we consider. We favored the block jackknife method for practical analysis due to implementation advantages: the block jackknife method can be implemented using the standard analysis software *PLINK*[7], while the cluster-based method currently cannot. Though the cluster-based method has lower theoretical time complexity, computing it in practice introduces data processing bottlenecks, making it significantly slower to execute on large-scale genomic data than the block jackknife.

#### S5.1 Close relatedness among families introduces confounding in uncertainty estimates

**Model setup.** Biobank-scale datasets rarely represent a random sample of independent families. The close relatedness among families violates the assumption of independence among sibships. To evaluate the performance of the variance estimators under such conditions, we simulated between-sibship covariance arising from extended family relatedness.

Specifically, we modeled a scenario in which the fathers of sequentially paired families (families $2k-1$ and $2k$, for $k = 1, \ldots, \frac{n_{sibships}}{2}$) were siblings, making the offspring of the two families first cousins. We modeled family-specific allelic effects as covarying between related families,

$$\text{Corr}[\beta_{2k-1}, \beta_{2k}] = 0.4, \quad k = 1, \ldots, \frac{n_{sibships}}{2}. \tag{26}$$

Notably, in the within-sibship difference regression, the genotypic covariance between the two related families cancels out. Because any sibling in family $i$ shares the exact same degree of genetic relatedness with any sibling in family $j$, all four pairwise genotypic covariances between the two families are equal:

$$\text{Cov}[X_{k,i}, X_{l,j}] = c, \ c \text{ is a constant, for any } k, l \in \{1, 2\},$$

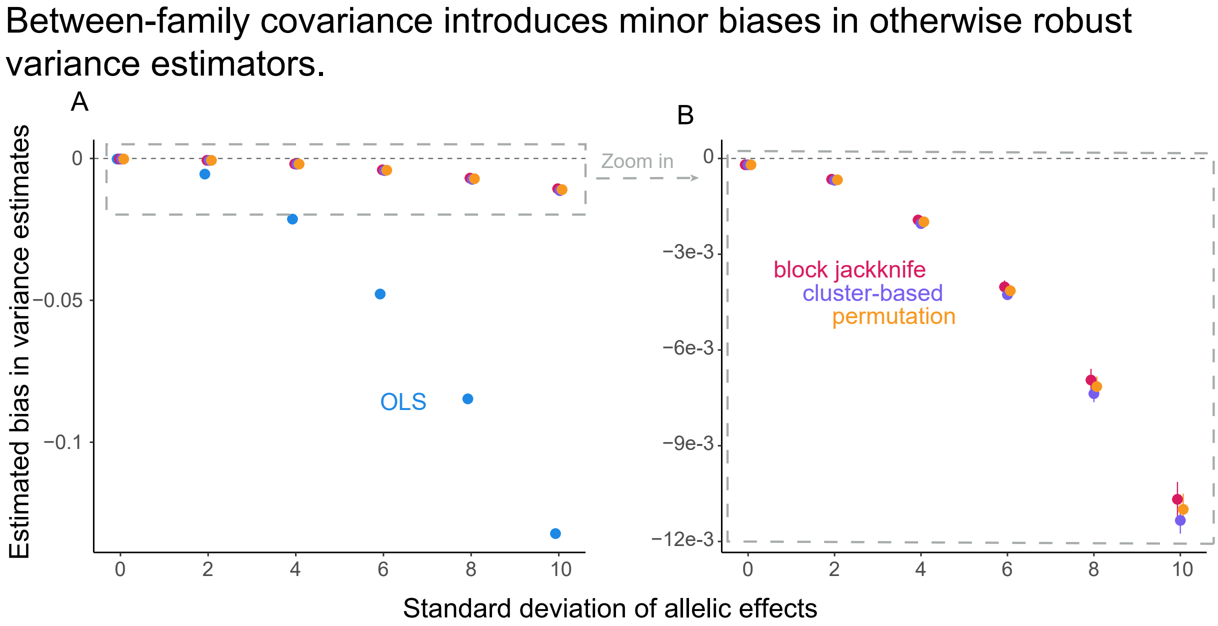

**Figure S4. Relatedness among families introduces biases in uncertainty estimates.** Markers show the estimated bias (**Eq. 28**) of the four variance estimators. Error bars show the mean ± 1.96 · SE of the estimated bias across 5,000 simulation replicates. In each replicate, we simulated 5,000 sibling pairs using a three-generation pedigree in which each pair of consecutive families ($2k-1$ and $2k$) is related: one parent from family $2k-1$ and one parent from family $2k$ are full siblings, making the offspring across the two families first cousins. Allelic effects between paired cousins were simulated with a correlation of $\text{corr}[\beta_{2k-1}, \beta_{2k}] = 0.4, k = 1, \ldots, \frac{n_{sibships}}{2}$, while the standard deviation $\tau$ was varied along the x-axis. Grandparental and unrelated parental genotypes at the focal SNP were drawn from Hardy-Weinberg equilibrium at a maf of 10%, with Mendelian segregation applied in both generations. The baseline allelic effect was set to zero. **(A)** The OLS method is heavily biased by heterogeneous allelic effects, consistent with our main text conclusions, whereas the block jackknife, permutation, and the cluster-based methods show a much smaller bias. **(B)** A zoomed-in view of the dashed rectangle in (A). The three other estimators (other than the OLS estimator) exhibit a subtle negative bias that grows in magnitude with the standard deviation of allelic effects.

the covariance between the within-family genotypic differences is:

$$
\begin{aligned}
\text{Cov}[\Delta\text{X}_\text{i}, \Delta\text{X}_\text{j}] &= \text{Cov}[X_{1,i} - X_{2,i}, X_{1,j} - X_{2,j}] \\
&= \text{Cov}[X_{1,i}, X_{1,j}] - \text{Cov}[X_{1,i}, X_{2,j}] - \text{Cov}[X_{2,i}, X_{1,j}] + \text{Cov}[X_{2,i}, X_{2,j}] \\
&= c - c - c + c \\
&= 0.
\end{aligned}
$$

This fact allows us to interpret changes in bias as stemming from the correlation in allelic effects rather than genotypic covariance.

**Simulations and results.** Based on the individual model (**Eq. 1** in the main text) and the covariance between sibships (**Eq. 26**), we simulated 5,000 sibling pairs with one focal SNP. To generate genotypes, we first simulated a three-generation pedigree. In the grandparent generation (G0), grandparental genotypes were drawn from the Hardy-Weinberg equilibrium with minor allele frequency 10%. Two first-generation (G1) siblings were then produced by Mendelian transmission from the same grandparental pair, making them full siblings to each other. Each G1 sibling mated with an unrelated spouse whose genotype was drawn independently from the population, and the resulting offspring constituted the focal sibling pairs (G2). Under this design, the focal siblings within each family are full siblings, and the focal siblings across paired families (families $2k-1$ and $2k$) are first cousins.

Allelic effects were drawn from a bivariate normal distribution that allowed for correlation between cousin-sibships:

$$\begin{pmatrix}\beta_{2k-1}\\ \beta_{2k}\end{pmatrix} \sim N\left(\begin{pmatrix}0\\0\end{pmatrix}, \begin{pmatrix}\tau^2 & 0.4\tau^2\\ 0.4\tau^2 & \tau^2\end{pmatrix}\right), \quad k = 1, \dots, \frac{n_{\text{sibships}}}{2},$$

so that the correlation between allelic effects in paired cousin families is fixed at 0.4 across all parameter settings, while the standard deviation of allelic effects ($\tau$) varies. The non-focal terms within each family were drawn from a bivariate normal distribution with variance $\sigma_0 = 4$ and within-family correlation $\rho_e = 0.06$, reflecting shared genetic background and environment between siblings.

We varied the standard deviation of allelic effects, taking $\tau \in \{0, 2, 4, 6, 8, 10\}$, with the baseline allelic effect fixed at zero and homoskedastic non-focal variation. For each value of $\tau$, we ran 5,000 simulation replicates. The true variance of the estimated allelic effects, $\text{Var}[\hat{\beta}]$, was estimated using the empirical (sample) variance among the $5,000$ simulations:

$$\widehat{\text{Var}}_{emp}[\hat{\beta}] = \frac{1}{5000-1}\sum_{sim=1}^{5000}(\hat{\beta}_{sim} - \bar{\hat{\beta}})^2, \tag{27}$$

where $\bar{\hat{\beta}} = \frac{1}{5000}\sum_{sim=1}^{5000}\hat{\beta}_{sim}$. We then evaluated each variance estimator by calculating its estimated bias:

$$\widehat{\text{Bias}}\left[\widehat{\text{Var}}_{method}[\hat{\beta}]\right] = \widehat{\text{Var}}_{method}[\hat{\beta}] - \widehat{\text{Var}}_{emp}[\hat{\beta}]. \tag{28}$$

Our results show that when the allelic effects are heterogeneous, the OLS method is substantially biased, consistent with our primary findings. In contrast, the block jackknife, cluster-based, and permutation methods show much lower bias (**Fig. S4A**). A closer inspection reveals that the three otherwise robust estimators exhibit a slight negative bias that increases alongside the standard deviation of the allelic effect (**Fig. S4B**). This negative bias is expected because none of these methods explicitly incorporates cross-sibship correlation; they treat cousin-sibships as independent units and therefore overestimate the effective sample size and underestimate uncertainty. Overall, these results demonstrate that while the block jackknife and cluster-based methods are far more robust than the OLS method, they are similarly affected by unmodeled between-sibship relatedness.

#### S5.2 Population structure can introduce bias in uncertainty estimates.

**Model setup.** Unmodeled population structure is suspected to be a major confounder in large-scale GWAS[8–11]. While sib-based allelic effect estimates are considered largely robust

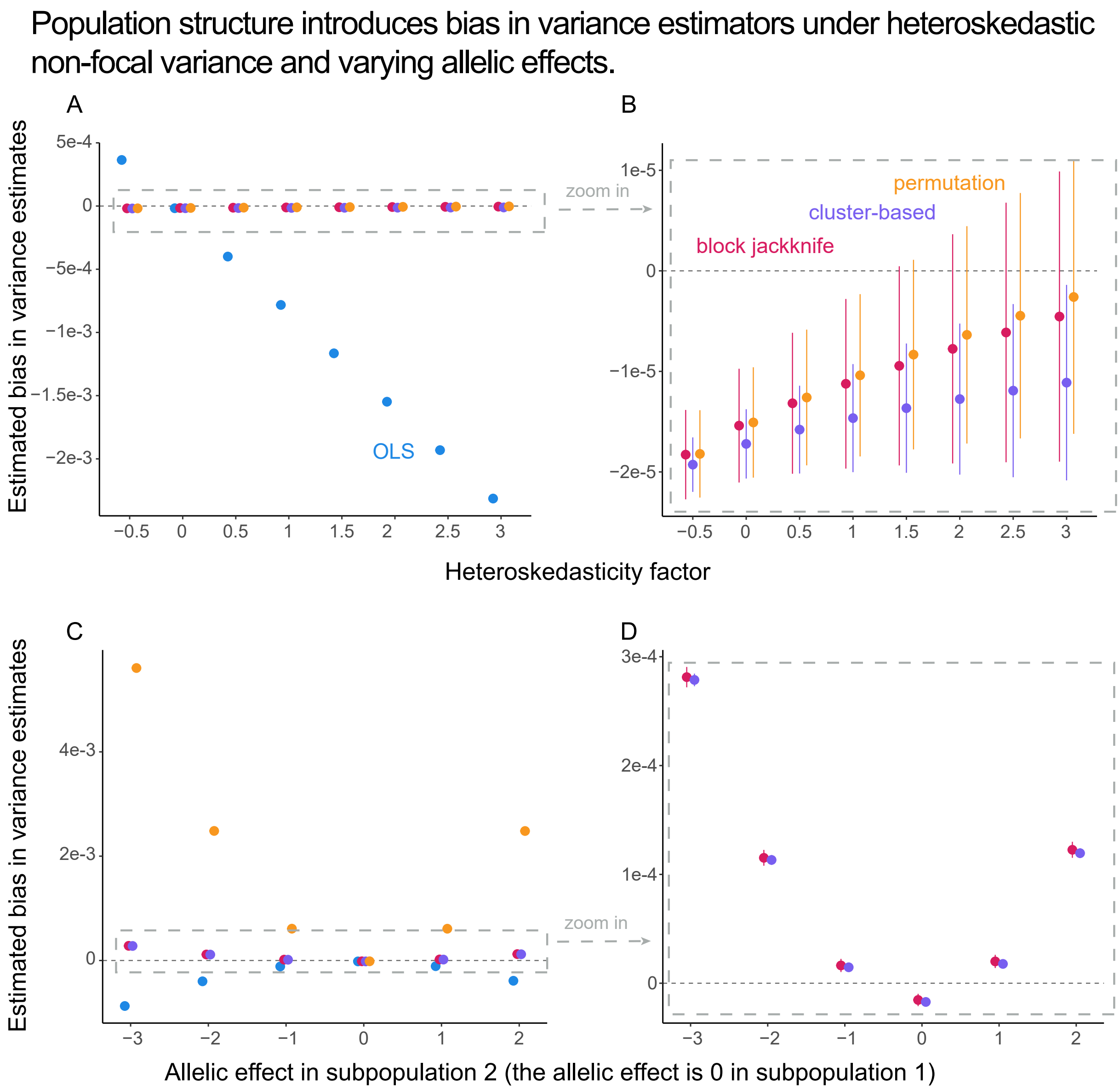

**Figure S5. Population structure introduces biases in uncertainty estimates.** Markers show the estimated bias (**Eq. 28**) of the four variance estimators. Error bars show the mean $\pm 1.96 \cdot \text{SE}$ of the estimated bias across 5,000 simulation replicates. In each replicate, we simulated 5,000 sibling pairs in which half come from subpopulation 1 with parental genotypes drawn from a Hardy-Weinberg equilibrium at a maf of 10% and the other half from subpopulation 2 at a maf of 40%. Unless otherwise specified, the baseline allelic effect and heteroskedasticity factor were set to zero. **(A-B)** vary the heteroskedasticity factor. **(A)** The OLS variance estimator exhibits massive bias, consistent with the main text conclusions. The other three methods are not severely influenced. **(B)** A zoomed-in view of the dashed rectangle in (A) reveals that the otherwise unbiased variance estimators still exhibit a slight bias that increases with the heteroskedasticity factor. **(C-D)** vary the baseline allelic effect in subpopulation 2 (the allelic effect is set to zero in subpopulation 1). **(C)** Both the OLS and permutation methods are heavily biased by heterogeneous non-zero allelic effects in the combined population. **(D)** A zoomed-in view of the dashed rectangle in (C) reveals that the block jackknife and cluster-based methods exhibit smaller bias when the allelic effects differ across the subpopulations.

to population structure, estimators of their variance may still be impacted. To evaluate the performance of the variance estimators in the presence of population structure, we simulated a cohort composed of two distinct subpopulations with differing minor allele frequencies and local linkage disequilibrium (LD) structures.

Specifically, we modeled a scenario in which the parent generation consisted of two subpopulations, each contributing 2500 families to the final combined cohort. Within each subpopulation, parental genotypes were drawn independently under the Hardy-Weinberg equilibrium with subpopulation-specific mafs (0.1 in subpopulation 1, 0.4 in subpopulation 2). We then performed sib-GWAS on the siblings from the combined population. Under this stratified framework, we evaluated the estimators under two scenarios:

(1) the non-focal variation is heteroskedastic;

(2) the true allelic effects differ across the subpopulations.

**Simulation and results: Heteroskedastic non-focal variation under population structure.** First, we assessed whether population structure exacerbated bias under heteroskedastic non-focal variation. Following the heteroskedastic assumption (**Eq. 4**) in the main text, we modeled the non-focal variance as a function of the non-focal genotype contrast:

$$\mathrm{Var}[\Delta\epsilon_i] = \sigma_0 + \sigma_1 \cdot \Delta X_i^2.$$

We varied the heteroskedasticity factor (equals $\sigma_1$ here, since $\tau^2 = 0$) $\sigma_1 \in \{-0.3, -0.2, \ldots, 0.3\}$, with the baseline allelic effect fixed at zero. For each value of $\sigma_1$, we ran 5,000 simulation replicates and estimated the bias of each variance estimator by **Eq. 28**.

Our results show that when the non-focal variance depends on the focal genotype contrast, the OLS method is substantially biased, consistent with our primary conclusions. In contrast, the block jackknife, cluster-based, and permutation methods exhibit much lower bias (**Fig. S5A**). A closer inspection reveals that the three otherwise robust estimators exhibit slight bias that increases with the heteroskedasticity factor (**Fig. S5B**). These results indicate that though the block jackknife and cluster-based methods are more robust than the OLS estimator, they remain similarly affected by unmodeled population structure.

**Simulation and results: subpopulation-specific allelic effects.** We next assessed whether population structure introduced additional challenges when the true allelic effect differed across subpopulations. Such differences can arise, for example, from distinct LD patterns across ancestral backgrounds, where a focal tag SNP's linkage to underlying causal variants differs among subpopulations. Specifically, we modeled the baseline allelic effect as 0 in subpopulation 1 and varied its value in subpopulation 2 within the set: $\{-3, -2, -1, 0, 1, 2\}$, with the heteroskedasticity factor fixed at zero. For each parameter setting, we ran 5,000 simulation replicates and estimated the bias of each variance estimator using **Eq. 28**.

Our results show that the permutation method is upward-biased in the presence of a non-zero allelic effect, while the OLS method is downward-biased because the allelic effect is heterogeneous in the combined population, consistent with our findings in the main text. In contrast, the block jackknife and cluster-based methods show much lower bias (**Fig. S5C**). A closer inspection reveals that the two otherwise robust estimators exhibit a slight bias that increases alongside the difference in the baseline allelic effects between subpopulations (**Fig. S5D**). These results demonstrate that while the block jackknife and cluster-based methods are more robust than the OLS and permutation methods, they are still challenged to a similar extent by unmodeled population structure.

#### S6 Generalization to the heterogeneous family sizes

Throughout the main text, we assumed two siblings per family and conducted within-sibship difference regression. However, data with more than two siblings per family are increasingly available. Recent work has leveraged multiple siblings per family and incorporated parental genotype imputation[5,12]. Here, we conduct simulations to show that our results in the main text generalize to settings with multiple siblings per family and specifically that the block jackknife method remains asymptotically unbiased.

**Model Setup.** We adopted the individual-level model from the main text (**Eq. 1**). The phenotype of each sibling was modeled as the sum of family-specific baseline ($Y_{0,i}$), the additive effect of the focal allele, and a non-focal term ($\epsilon_{j,i}$). The allelic effect was allowed to vary across sibships:

$$\beta_i = \beta_0 + U_i, \quad U_i \overset{i.i.d}{\sim} (0, \tau^2).$$

To accommodate multiple siblings per family, we followed the additive model in Howe et al.[5]:

$$Y_{j,i} = \alpha_0 + \beta_W (X_{j,i} - \bar{X}_i) + \beta_B \bar{X}_i + \epsilon_{j,i}, \tag{29}$$

where $n_{sibs,i}$ denotes the number of siblings in sibship $i$, and $\bar{X}_i = \frac{\sum_{j=1}^{n_{sibs,i}} X_{j,i}}{n_{sibs,i}}$ is the mean genotype within sibship $i$. The within-sibship effect $\beta_W$ corresponds to the allelic effect $\beta_i$ in **Eq. 3**. To incorporate heteroskedastic non-focal variation, we extended **Eq. 4** to the multiple-sibling setting:

$$\mathrm{Var}[\epsilon_{j,i}] = \sigma_0 + \sigma_1 \cdot \sum_{j=1}^{n_{sibs,i}} (X_{j,i} - \bar{X}_i)^2.$$

As in the two-sibling setting, the heteroskedasticity factor is defined as $\mathrm{Var}[\beta_i] + \sigma_1$.

**The simulation procedure.** We simulated one focal SNP in a sample of 5,000 families, where 2,500 families had 2 siblings, 1,500 families had 3 siblings, and 1,000 families had 4 siblings. Parental genotypes were drawn under the Hardy-Weinberg equilibrium at a minor allele frequency of 10%. Each sibling's genotype was obtained by independent Mendelian transmission from the two parents. We varied three parameters separately, holding the others at their defaults (baseline allelic effect $\beta_0 = 0$, standard deviation of the allelic effects $\tau = 0$, and the non-focal variation does not depend on the focal genotype contrast $\sigma_1 = 0$): heteroskedasticity factor ($\sigma_1 \in \{-0.3, -0.2, \ldots, 0.3\}$), the baseline allelic effect ($\beta_0 \in \{-2, -1, 0, 1, 2\}$), and its standard deviation ($\tau = \sqrt{\mathrm{Var}[\beta_i]} \in \{0, 2, 4, 6, 8\}$). For each parameter setting, we ran 5,000 simulation replicates and estimated the bias of each variance estimator using **Eq. 28**.

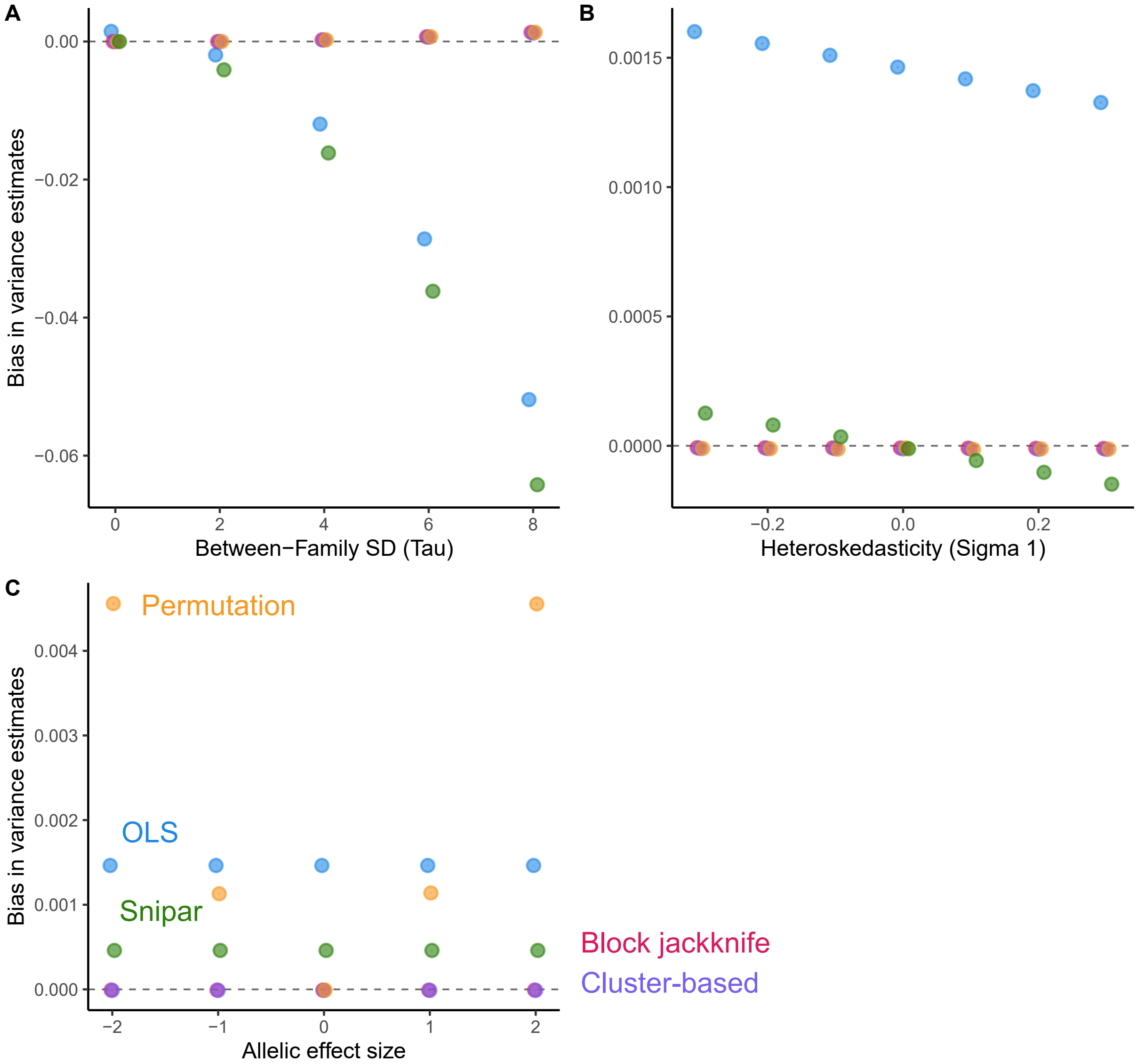

**Figure S6. Main text conclusions generalize to the heterogeneous family sizes setting.** Markers show the estimated bias of each variance estimator. Error bars show the mean ± 1.96 · SE of the estimated bias among the 5,000 simulation replicates. In each replicate, we modeled 5,000 families, with 2,500 families having 2 siblings, 1,500 families having 3 siblings, and 1,000 families having 4 siblings. The parental genotypes were sampled from the Hardy-Weinberg equilibrium at a maf of 10%. By default, the baseline allelic effect and the heteroskedasticity factor were set to 0. **(A)** varies the standard deviation of the allelic effects. **(B)** varies the heteroskedasticity factor. **(C)** varies the baseline effect.

**The block jackknife method remains asymptotically unbiased.** Our simulations show that the block jackknife estimator remains asymptotically unbiased in the three-sibling setting under the scenarios we consider (**Fig. S6**), consistent with our findings in the main text. The OLS variance estimates and the estimates from snipar are biased in the presence of heterogeneous allelic effects and heteroskedastic non-focal variation (**Fig. S6(A-B)**), consistent with the two-sibling setting. The permutation method is biased in the presence of a non-zero baseline allelic effect (**Fig. S6C**), consistent with the conclusion for the 2-sibling scenario. (**Fig. S6(B-C)**), consistent with the two-sibling setting. The permutation method is biased in the presence of a non-zero baseline allelic effect (**Fig. S6A**), consistent with the conclusion for the 2 siblings scenario.

### S7 Additional Supplementary Figures

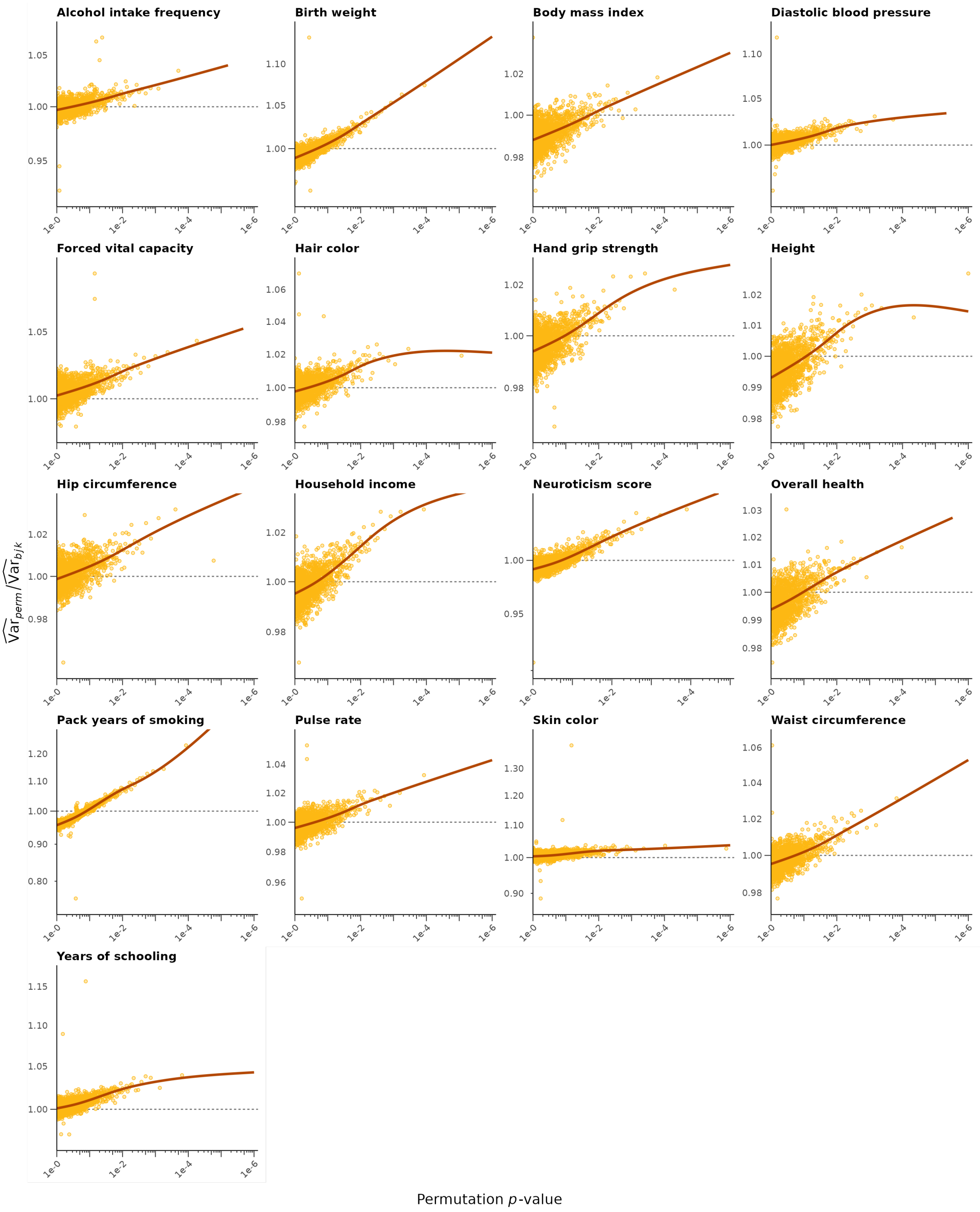

**Figure S7. Permutation variance estimator bias across all 17 traits from the UKB.** Each panel shows the results for one trait. Each point represents the mean $\log_2$ ratio of the permutation to block jackknife variance estimates across 300 SNPs binned by the permutation p-value of the allelic effect estimate. Lines show cubic splines fitted over the raw unbinned ratios. Consistent with the theoretical prediction in **Eq. 8**, the permutation estimator increasingly overestimates variance as evidence for nonzero allelic effect strengthens (smaller p-values) across all traits.

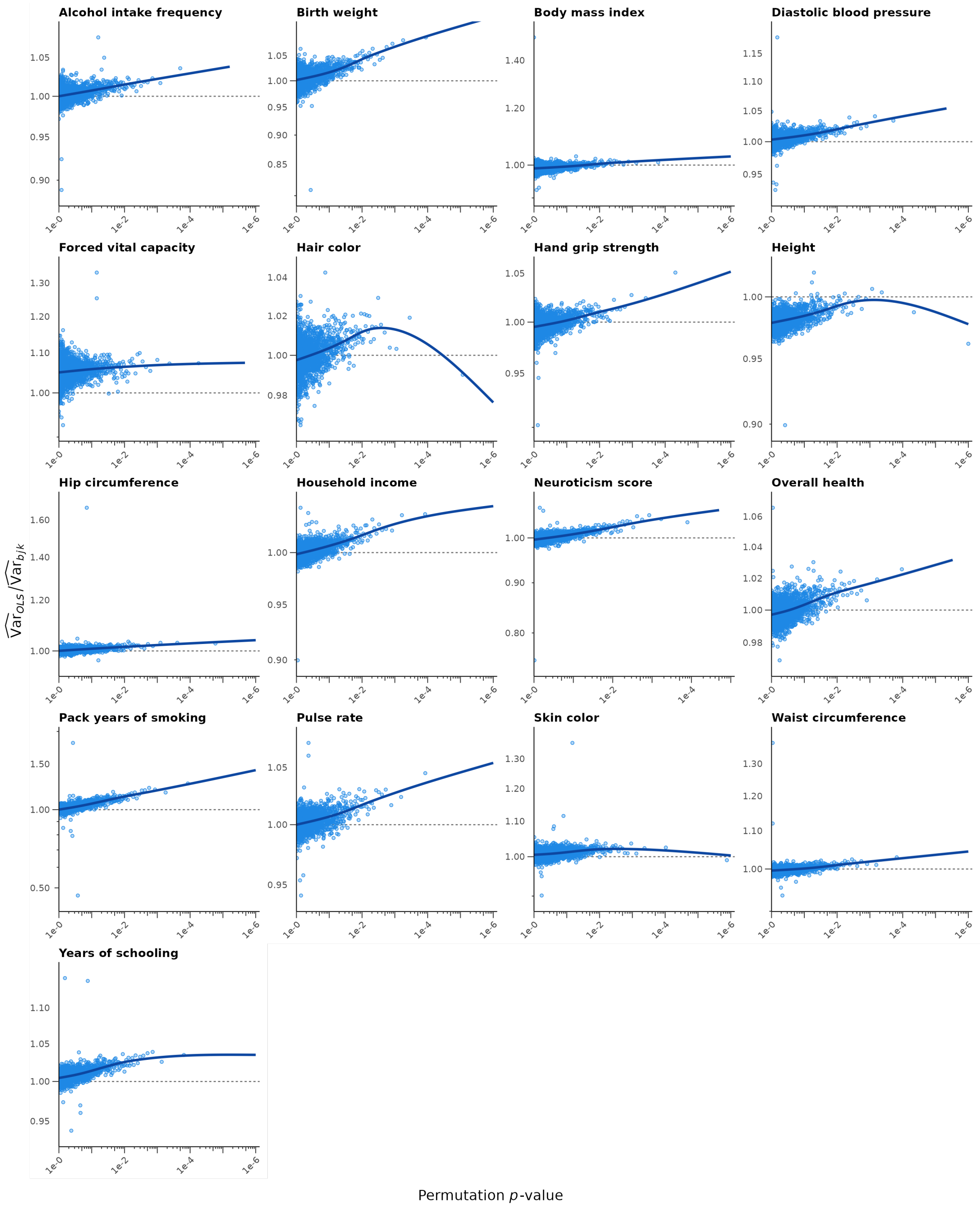

**Figure S8. OLS variance estimator bias across all 17 traits from the UKB.** Each panel shows the results for one trait. Each point represents the mean $\log_2$ ratio of the OLS to block jackknife variance estimates across 300 SNPs binned by the permutation p-value of the allelic effect estimate. Lines show cubic splines fitted over the raw unbinned ratios. Consistent with the theoretical prediction in **Eq. 7**, the OLS estimator increasingly deviates from the block jackknife benchmark as evidence for nonzero allelic effects strengthens (smaller p-values), consistent across all traits.

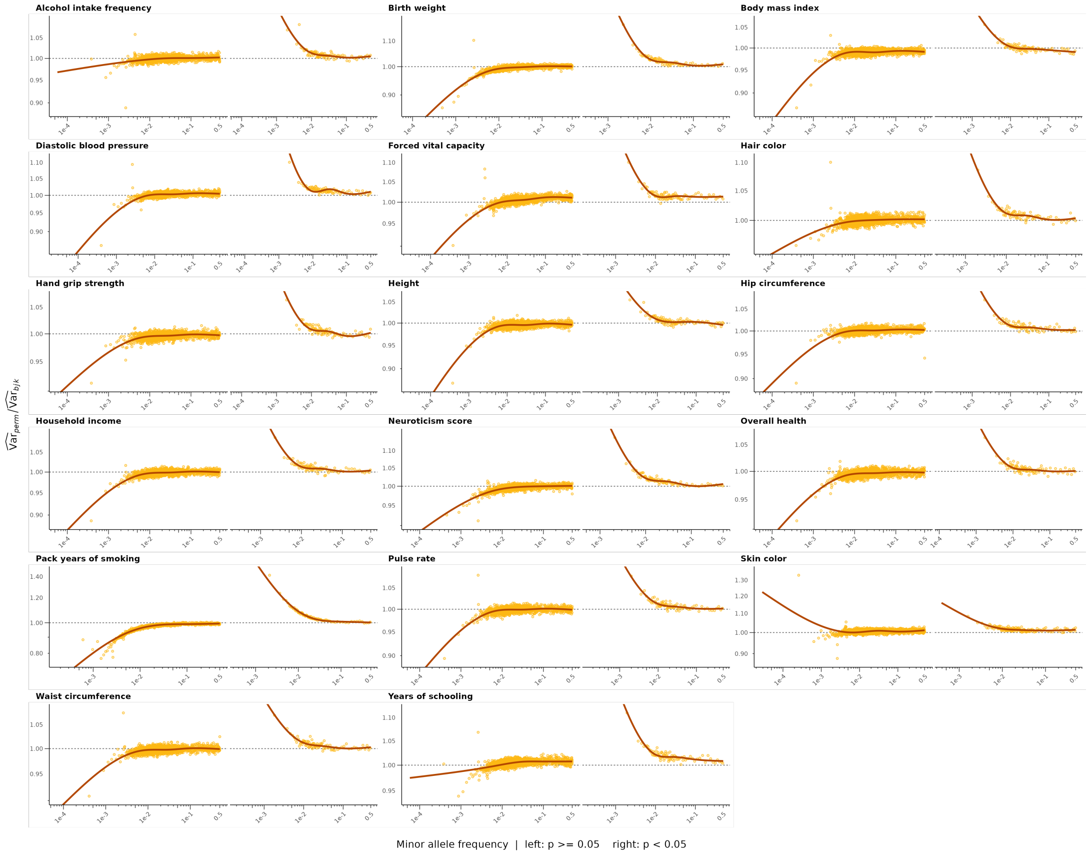

**Figure S9. Permutation variance estimator bias across all 17 traits from the UKB.** Each panel shows the results for one trait, split by whether the permutation p-value of the allelic effect was below 0.05 or at least 0.05. Each point represents the mean $\log_2$ ratio of the permutation to block jackknife variance estimates across 300 SNPs binned by minor allele frequency. Lines show cubic splines fitted over the raw unbinned ratios. Consistent with the theoretical prediction in **Eq. 8**, the magnitude of the permutation bias is larger for rarer alleles across all traits.

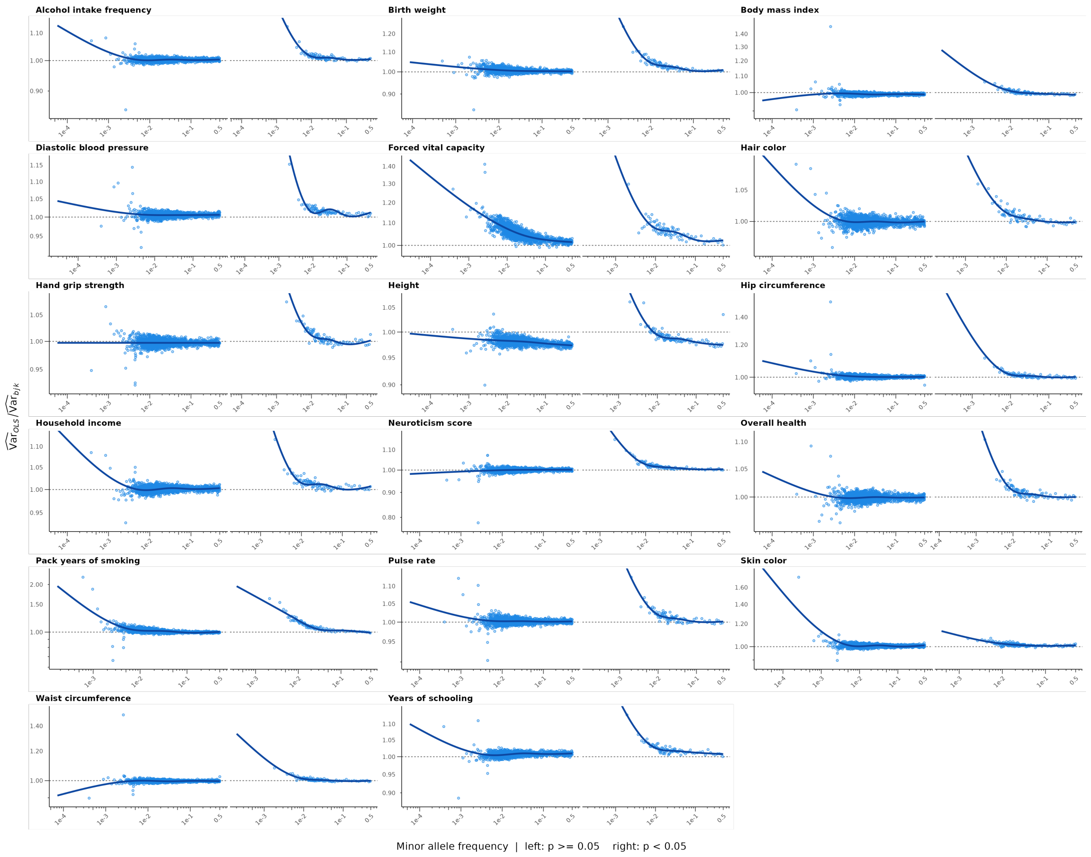

**Figure S10. OLS variance estimator bias across all 17 traits from the UKB.** Each panel shows the results for one trait, split by whether the permutation p-value of the allelic effect was below 0.05 or at least 0.05. Each point represents the mean $\log_2$ ratio of the OLS to block jackknife variance estimates across 300 SNPs binned by minor allele frequency. Lines show cubic splines fitted over the raw unbinned ratios. Consistent with the theoretical prediction in **Eq. 7**, the magnitude of the OLS bias is larger for rarer alleles across all traits.

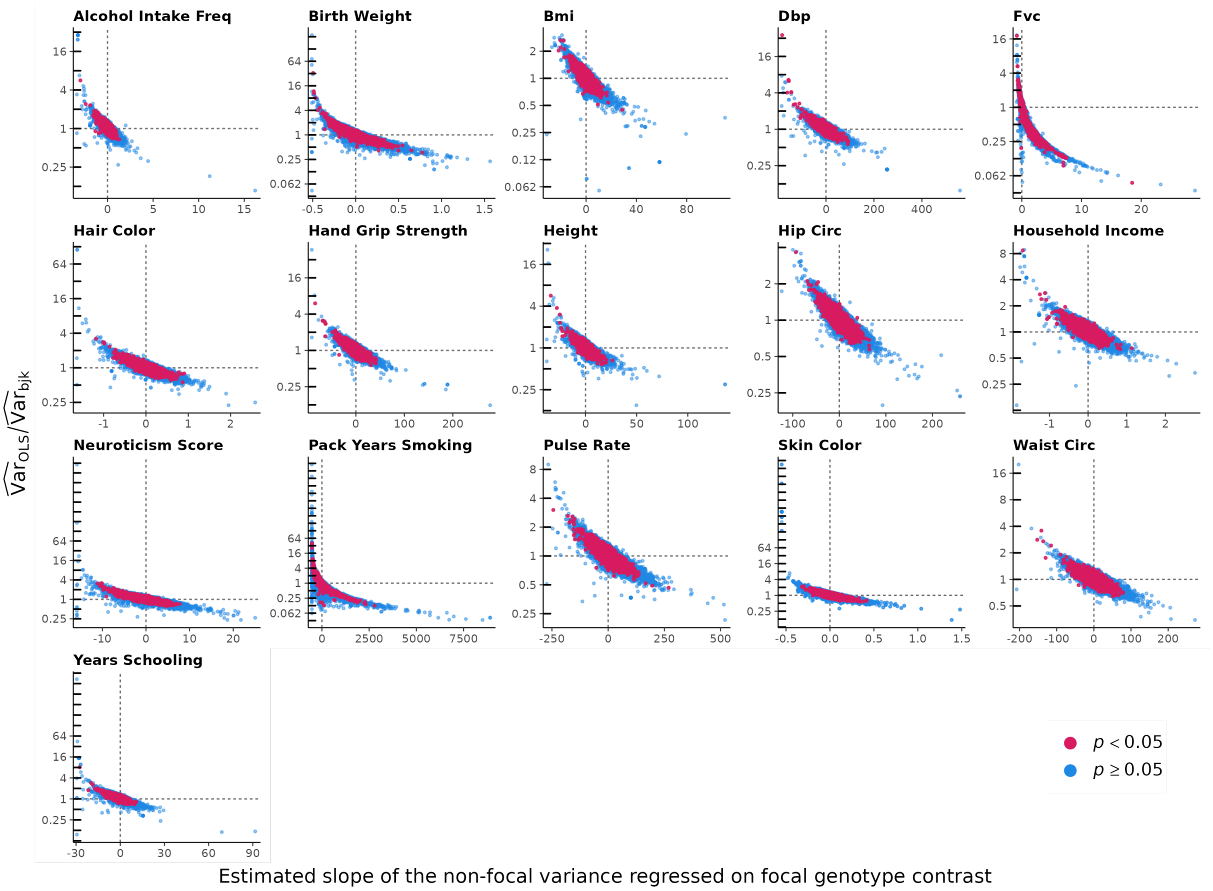

**Figure S11. Heteroskedasticity factor drives the direction of the bias in the OLS variance estimator.** Each panel shows results for one trait. The estimated slope $\hat{\sigma}_1$ describes how non-focal variance changes with focal genotype contrast (**Eq. 4**); the $\log_2$ ratio of the OLS to the block jackknife variance estimate shows the magnitude that the OLS estimator deviates from the block jackknife benchmark. Consistent with the theoretical prediction in **Eq. 7**, the OLS variance estimate falls below the block jackknife benchmark for SNPs with positive $\sigma_1$ and rises above it for SNPs with negative $\sigma_1$ across all traits.

### S8 Supplementary Table

| Trait | Sib-GWAS Sample Size (Sibling Pairs) | UK Biobank Field ID |
|---|---|---|
| Alcohol intake frequency | 17,328 | 1558 |
| Birth weight | 6,569 | 20022 |
| BMI | 17,257 | 21001 |
| Diastolic blood pressure | 14,717 | 4079, 6153, 6177 |
| Forced vital capacity | 14,522 | 3062 |
| Hair color | 17,292 | 1747 |
| Hand grip strength | 17,178 | 46, 47 |
| Height | 17,284 | 50 |
| Hip circumference | 17,297 | 49 |
| Household income | 13,125 | 738 |
| Neuroticism score | 8,358 | 20127 |
| Overall health rating | 17,234 | 2178 |
| Pack years of smoking | 2,163 | 20161 |
| Pulse rate | 15,415 | 102 |
| Skin color | 16,924 | 1717 |
| Waist circumference | 17,301 | 48 |
| Years of schooling | 12,076 | 6138 |

**Table S1. Details of traits analyzed in the UK Biobank.** Sample sizes reflect the number of White British full sibling pairs with complete phenotypic data for each trait, after removing individuals identified as part of a sibling pair from the standard-GWAS cohort.

### References

[1] Yuval B. Simons, Kevin Bullaughey, Richard R. Hudson, and Guy Sella. "A population genetic interpretation of GWAS findings for human quantitative traits". *PLOS Biology* 16.3 (Mar. 2018). Ed. by Greg Gibson, e2002985. ISSN: 1545-7885. DOI: 10.1371/journal.pbio.2002985.

[2] E. Koch et al. "Genetic association data are broadly consistent with stabilizing selection shaping human common diseases and traits" (2024). DOI: 10.1101/2024.06.19.599789.

[3] Roshni A Patel et al. "Characterizing selection on complex traits through conditional frequency spectra". *Genetics* 229.4 (Dec. 2024). Ed. by S Gravel. ISSN: 1943-2631. DOI: 10.1093/genetics/iyae210.

[4] George Casella and Roger Berger. *Statistical inference.* Chapman and Hall/CRC, 2024.

[5] Laurence J. Howe et al. "Within-sibship genome-wide association analyses decrease bias in estimates of direct genetic effects". *Nature Genetics* 54.5 (May 2022), pp. 581–592. ISSN: 1546-1718. DOI: 10.1038/s41588-022-01062-7.

[6] Halbert White. "A Heteroskedasticity-Consistent Covariance Matrix Estimator and a Direct Test for Heteroskedasticity". *Econometrica* 48 (1980), pp. 817–838. ISSN: 0012-9682. DOI: 10.2307/1912934. URL: https://www.jstor.org/stable/1912934.

[7] Shaun Purcell et al. "PLINK: A Tool Set for Whole-Genome Association and Population-Based Linkage Analyses". *The American Journal of Human Genetics* 81.3 (2007), pp. 559–575. ISSN: 0002-9297. DOI: 10.1086/519795.

[8] Samuel Pattillo Smith et al. "A Litmus Test for Confounding in Polygenic Scores" (Feb. 2025). DOI: 10.1101/2025.02.01.635985.

[9] Nick Barton, Joachim Hermisson, and Magnus Nordborg. "Why structure matters". *eLife* 8 (Mar. 2019). ISSN: 2050-084X. DOI: 10.7554/eLife.45380.

[10] Alexander I. Young, Stefania Benonisdottir, Molly Przeworski, and Augustine Kong. "Deconstructing the sources of genotype-phenotype associations in humans". *Science* 365.6460 (2019), pp. 1396–1400. ISSN: 1095-9203. DOI: 10.1126/science.aax3710.

[11] Guy Sella and Nicholas H. Barton. "Thinking About the Evolution of Complex Traits in the Era of Genome-Wide Association Studies". *Annual Review of Genomics and Human Genetics* 20.1 (Aug. 2019), pp. 461–493. ISSN: 1545-293X. DOI: 10.1146/annurev-genom-083115-022316.

[12] Alexander I. Young et al. "Mendelian imputation of parental genotypes improves estimates of direct genetic effects". *Nature Genetics* 54.6 (2022), pp. 897–905. ISSN: 1546-1718. DOI: 10.1038/s41588-022-01085-0.
